# Human-specific gene expansions contribute to brain evolution

**DOI:** 10.1101/2024.09.26.615256

**Authors:** Daniela C. Soto, José M. Uribe-Salazar, Gulhan Kaya, Ricardo Valdarrago, Aarthi Sekar, Nicholas K. Haghani, Keiko Hino, Gabriana La, Natasha Ann F. Mariano, Cole Ingamells, Aidan Baraban, Zoeb Jamal, Tychele N. Turner, Eric D. Green, Sergi Simó, Gerald Quon, Aida M. Andrés, Megan Y. Dennis

## Abstract

Duplicated genes expanded in the human lineage likely contributed to brain evolution, yet challenges exist in their discovery due to sequence-assembly errors. We used a complete telomere-to-telomere genome sequence to identify 213 human-specific gene families. From these, 362 paralogs were found in all modern human genomes tested and brain transcriptomes, making them top candidates contributing to human-universal brain features. Choosing a subset of paralogs, long-read DNA sequencing of hundreds of modern humans revealed previously hidden signatures of selection, including for T-cell marker *CD8B*. To understand roles in brain development, we generated zebrafish CRISPR “knockout” models of nine orthologs and introduced mRNA-encoding paralogs, effectively “humanizing” larvae. Our findings implicate two genes in possibly contributing to hallmark features of the human brain: *GPR89B* in dosage-mediated brain expansion and *FRMPD2B* in altered synapse signaling. Our holistic approach provides insights and a comprehensive resource for studying gene expansion drivers of human brain evolution.

## Introduction

Significant phenotypic features distinguish modern humans from closely related great apes^1–4^. Arguably, one of the most compelling innovations relates to changes in neuroanatomy, including an expanded neocortex and increased complexity of neuronal connections, which allowed the development of novel cognitive features such as reading and language^5^. While previous work implicated human-specific single-nucleotide variants (SNVs) that impact genes leading to altered brain features, including *FOXP2*^6,7^ and human-accelerated regions^8^, a majority of top gene candidates are the result of segmental duplications (SDs; genomic regions >1 kbp in length that share high sequence identity [>90%])^9–11^. SDs can give rise to new gene paralogs with the same function, altered functions, or that antagonize conserved, ancestral paralogs^12^ and contribute more to genetic divergence across species than SNVs^13^. Previous comparisons of great ape genomes have identified >30 human-specific gene families and hundreds of paralogs important in neurodevelopment and enriched at genomic hotspots associated with neuropsychiatric disorders^14–16^. Of these, a handful of genes have been found to function in brain development using model systems, including *SRGAP2C*^17,18^, *NOTCH2NL*^19–21^, *ARHGAP11B*^22–24^, *TBC1D3*^25^ *CROCCP2*^26^, and *LRRC37B*^27^. Most studies have leveraged mice to study gene functions with recent studies expanding to cortical organoids, ferrets, and primates^28^. Despite their clear importance in contributing to neural features, most duplicate genes remain functionally uncharacterized due to the arduous nature of using such models.

SDs have largely eluded analyses because of difficulties in accurate genome assembly^29^ and discovering variants across nearly identical paralogs^30–34^. As such, many human-duplicated genes are likely left to be discovered. The telomere-to-telomere (T2T) human reference genome T2T-CHM13^35^, representing a gapless sequence of all autosomes and chromosome X, has enabled a more complete picture of SDs^36^ by incorporating hundreds of megabases missing from the previous human reference genome (GRCh38).

This new assembly corrects >8 Mbp of collapsed duplications^37^, including previously missing paralogs of human-specific duplicated gene families^14^ *GPRIN2*^36^ and *DUSP22*^37^. Here, using this new T2T genome, we identified thousands of recent gene duplications among hominids. By comparing great ape genomic data, we narrowed in on a set of paralogs unique within and fixed across modern humans. Transcriptomic datasets from the human brain identified genes most likely to contribute to neurodevelopment and function, providing a catalog of the candidate human-specific gene families contributing to brain evolution for further functional testing in model systems. Finally, we prioritized a set of duplicate gene families to characterize in more detail using long-read sequencing and systematic analysis in zebrafish to elucidate brain functions.

## Results

### Genetic analysis of human-duplicated genes

#### Identification of human gene duplications in T2T-CHM13

Understanding that highly identical SDs are enriched for human-specific duplications, we narrowed in on 97.8 Mbp of autosomal sequences sharing >98% identity with other genomic regions (or SD98) in the human T2T-CHM13^36,38^ (Figure 1A). These loci represent genes duplicated only in human lineage^14,15^ as well as expansions of duplicated gene families present in other great apes. Consistent with the notion that gene duplication can lead to functional innovation, a number of paralogs in the latter category have experienced recent changes along the *Homo* lineage in expression (e.g., *LRRC37B*^27^) or sequence content (e.g., *NOTCH2NL*, via interlocus gene conversion^19^). Focusing our analysis on autosomes, we identified 698 protein-encoding genes and 1,095 unprocessed pseudogenes representing possible mis-annotations of true protein-encoding genes^39^ (Table S1A;, 478 paralogs on sex chromosomes in Table S1B). This list includes well-known genes important in neurodevelopment (*SRGAP2C*, *ARHGAP11B*), disease (*SMN1* and *SMN2*^40^, *KANSL1*^41^), and adaptation (amylase^42–44)^, with 668 (37%) residing in previously missing or erroneous regions in GRCh38. Sequence read depth^36^ in modern humans (Simons Genome Diversity Project [SGDP], n=269^45^) verified that all paralogs had >2 gene-family diploid copy number (famCN; STAR Methods, Table S1C, Figure 1B).

**Figure 1.**
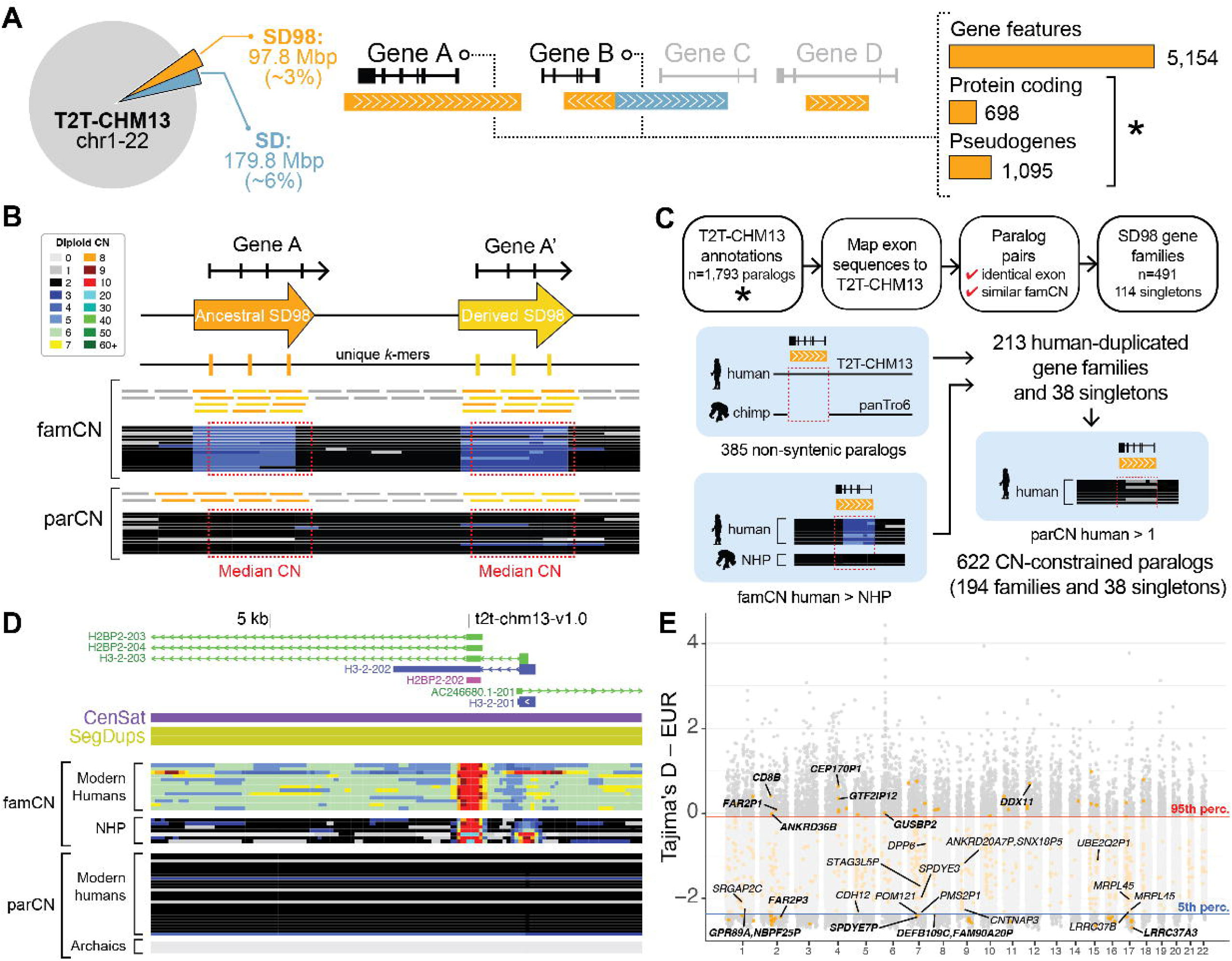
Genetic analysis of human-duplicated genes. (**A**) Diagram of segmental duplications (SDs; blue) and subset with >98% identity (SD98; orange) in T2T-CHM13 autosomes, including total number of nucleotides (Mbp) and genes overlapping SD98 regions. (**B**) Copy number (CN) estimation methods, including gene-family CN (famCN) and paralog-specific CN (parCN). Horizontal lines represent short reads mapping to unique (gray) and duplicated regions (orange and yellow). Heatmaps indicate CN estimates. (**C**) Pipeline for clustering and stratification of SD98 genes based on synteny with the chimpanzee reference and famCN comparisons between human and nonhuman primates (NHPs) (left). CN-constrained (fixed or nearly fixed) genes were flagged based on parCN values across human populations (right). (**D**) UCSC Genome Browser snapshot including gene models, centromeric satellites (CenSat), SDs (SegDup), and famCN and parCN predictions across sequenced individuals. (**E**) Distribution of Tajima’s D values (y-axis) from 1KGP individuals of European (EUR) ancestry genome wide (gray) and SD98 (orange) across human autosomal chromosomes (x-axis). SD98 windows above the 95th (red line) or below the 5th (blue line) percentiles are considered outlier D values (STAR Methods). All human-duplicated gene names with outlier D values in at least one tested ancestry are labeled. Also see Table S1 and Figure S1.

Based on sequence and famCN similarity, we clustered 1,679 of the paralogs into 491 multigene families, with many having 2–3 members (n=271 families) (Figures 1C and S1A,B). Three extreme high-copy gene families had >50 paralogs, including macrosatellite-associated *DUX4* and *DUB/USP17* as well as primate-specific *FAM90A*^46^. The remaining 114 paralogs were defined as “singletons” (Table S1C), with some failing to cluster due to high and variable copy numbers (CNs) (e.g., *CROCC* and *CROCCP2*) or only a small portion of the gene duplicated (e.g., *AIDA* and *LUZP2*). We defined 385 of these genes as human-specific, falling within non-syntenic human and chimpanzee reference regions^36^ (Table S1C, Figure 1C). Because several known human-specific paralogs were absent from our list (e.g., *NPY4R2*, *ROCKP1*, and *SERF1B*^14^), we also narrowed in on 97 human-expanded gene families and 27 singletons with higher famCN in humans versus nonhuman great apes (see STAR Methods). In total, we conservatively predict 213 gene families and 38 singletons comprising at least one human-specific duplicate paralog (Table S1D). Moving forward, we refer to any of the 1,002 paralogs within one of these gene families as a human-specific duplicated gene.

#### Variation of duplicated genes in modern humans

Positing that all humans should carry a functional version of a gene if important for a species-universal trait, we used *k*-mer-based paralog-specific copy number (parCN) estimates^47^ to identify 622 “CN constrained” genes (parCN≥0.5; >98% 1000 Genomes Project, 1KGP; n=2,504) and 125 paralogs “fixed” in humans (parCN∼2) (Table S1E). Thirteen genes represent *Homo sapiens*-specific gene duplications largely absent from four archaic human genomes^48–50^; these include *H3-2/H2BP2*, a member of a core *H2B* histone family involved in the structure of eukaryotic chromatin^51^, homologous with human-specific *H2BP1* and the ancestral *H2BC18* paralog (Figure 1D), as well as *FCGR1CP*, encoding an immunoglobulin gamma Fc Gamma Receptor implicated in regulating immune response^52^.

We identified 13 protein-encoding genes as loss-of-function intolerant using SNV data from hundreds of thousands of humans from gnomAD^53^ (Table S1A, Figure S1C), showing that deleterious mutations of these genes are depleted in human populations. The gnomAD (v3) metrics rely on variants identified in protein-encoding genes using the human reference genome hg19, which has known errors across SDs^54^ and misannotated pseudogenes. As such, all unprocessed pseudogenes and 32% of protein-encoding SD98 genes lacked gnomAD pLI and LOEUF scores. To circumvent these issues, we assessed SNV genetic diversity in the 1KGP cohort by Tajima’s D^37,55,56^ (Figures S1D–G) Genic SD98 loci exhibit significantly reduced Tajima’s D compared with non-coding regions, indicating increased functional constraint and matching results in non-duplicated regions (Figure S1H). We identified 15 CN-constrained human-duplicated genes with the most negative D values (<5^th^ percentile; see Methods) considered outliers, suggestive of signatures of positive or strong levels of purifying selection (Table S1F, Figures 1E and S1I). These included human-specific paralog *SRGAP2C* previously implicated in cortical neuronal migration and synaptogenesis^17,18^ as well as the uncharacterized *LRRC37A3* and the hominid-specific *LRRC37B*, recently found to function in cortical pyramidal neurons by impacting synaptic excitability^27^. We also identified nine genes exhibiting the highest D values (>95^th^ percentile), suggestive of signatures of balancing selection, including T-cell antigen *CD8B.* We note that duplicated genes exhibiting non-extreme Tajima’s D can also be functionally constrained and provide these results across all short-read accessible regions as a resource^57^. Collectively, variants discovered using the new T2T-CHM13 genome enabled the identification of human-duplicated genes potentially contributing to traits and diseases not previously assayed in genome-wide selection screens.

### Human-duplicated genes implicated in brain development

#### Connecting genetic variation of duplicated genes with neural traits

Considering gene ontology (GO) of paralogs from human-duplicated families (n=1,002) and CN constrained (n=622), we did not identify any enrichments of functional features. We note that only 24% (236/1,002) of duplicated genes can be assigned a GO versus 79% of all protein-encoding and unprocessed pseudogenes, highlighting that most paralogs have unknown functions. To narrow in on human-duplicate gene families contributing to neurocognitive features, we identified 341 paralogs (187 CN-constrained) with putative associations with brain-related phenotypes by intersecting with the genome-wide association studies (GWAS) catalog and UK Biobank^58^ (Table S1A, Figure S1J, STAR Methods). Many human-duplicated genes reside at genomic hotspots (n=305), such as *GPR89* paralogs at chromosome 1q21.1 with recurrent ∼2 Mbp deletions/duplications associated with autism and impacting brain size^59^. To better delineate copy-number variants (CNVs), we assayed parCN in an autism cohort (Simons Simplex Collection [SSC]; n=2,459 quad families ^60,61^). Eighteen duplicated genes residing at autism-associated hotspots, including chromosomes 15q25.2 (OMIM:614294) and 3q29 (OMIM:609425), show significant parCN differences in probands versus unaffected siblings (Wilcoxon signed-rank test, *q*-value<0.05) (Figure S1K). *De novo* CNVs impact 22 human-duplicated genes in autistic probands in contrast to six events impacting five paralogs in unaffected siblings (Fisher’s exact test, *p*-value=4.5×10^-4^) (Table S1G, Figure S1L). While a majority of impacted genes reside at known hotspots, some did not, including *CD8B2*, *FCGR1B*, *HYDIN* and *LIMS1*, representing possible contributors to autism.

#### Duplicated gene expression in the developing human brain

Nearly half of human-duplicated gene paralogs (455/1,002) are expressed during brain development (TPM≥1)^22,62–65^ (Table S1A, Figures 2A and S2A). While this represents a depletion versus the complete transcriptome (18,918/23,395; hypergeometric test, *p*-value=4.76×10^-146^), the number of brain-expressed genes increases to 58% for CN-constrained (1.3-fold enrichment over all human-duplicated genes, *p*-value=2.5×10^-24^) and to 84% for CN-constrained protein-encoding (1.4-fold enrichment, *p*-value=7.8×10^-30^) (Figures 2B and C). Similar expression increases are also observed in lymphoblastoid cell lines^66^ (Figure S2B) showing this is not specific to the brain but, instead, suggests true functional candidate genes are more likely to exist in CN-constrained protein-encoding genes.

**Figure 2.**
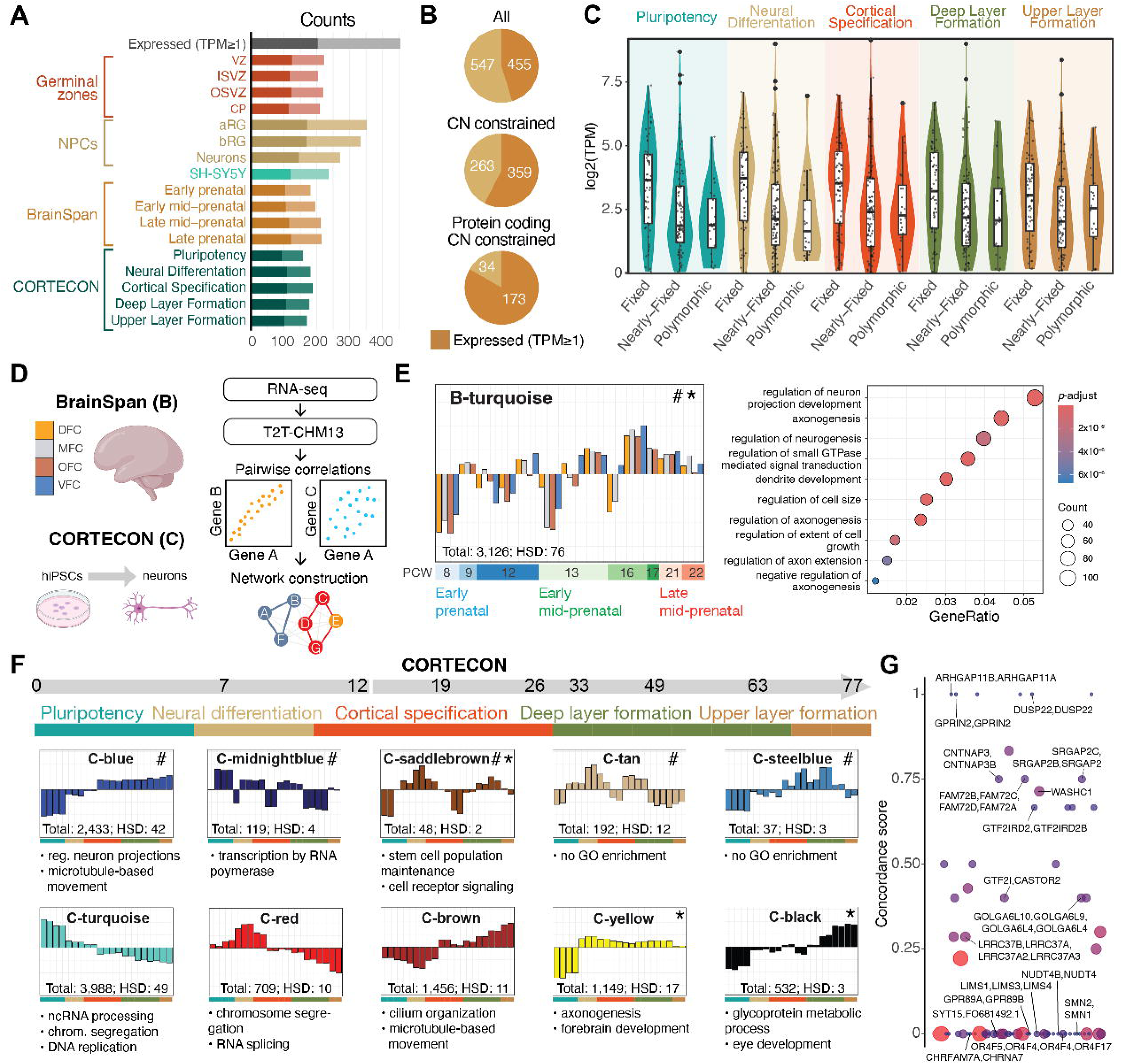
Duplicated gene expression in the developing human brain. (**A**) Counts of human-duplicated genes with transcripts per million (TPM) >1 in fetal brain datasets including germinal zones (VZ: ventricular zone, ISVZ: inner subventricular zone, OSVZ: outer subventricular zone, CP: cortical plate), neuronal progenitor cells (NPCs) (aRGs: apical radial glia, bRGs: basal radial glia), neuroblastoma cell line (SH-SY5Y), BrainSpan, and CORTECON. Protein-encoding genes are represented in darker shades. (**B**) Counts of expressed (dark orange) and non-expressed (light orange) human-duplicated genes across gene categories. (**C**) Human-duplicated gene expression in the CORTECON dataset stratified by copy number (CN). (**D**) Pipeline used for the weighted gene co-expression analysis (WGCNA). (**E**) The BrainSpan B-turquoise module, exhibiting an enrichment of human-duplicated genes (#) and autism-associated genes (*) plotted over developmental time (post-conception weeks, PCW) and bar colors representing brain regions (see **D**). Gene-ontology (GO) terms overrepresented among the co-expressed B-turquoise genes are depicted on the right. (**F**) Selected CORTECON WGCNA modules with enrichments (see **E**) and overrepresented GO terms indicated below. (**G**) CORTECON module assignment concordance scores are shown on the vertical axis for human-duplicated gene families. The size of each point corresponds to the number of members in the respective gene family. Also see Table S2, Figure S2, and Data S1.

We inferred possible functions of human-duplicated genes expressed during brain development using weighted gene co-expression network analysis (WGCNA)^67^. Assessment of post-mortem human prenatal frontal cortex transcriptomes spanning post-conception weeks 8 to 22 from BrainSpan^65^ revealed 17 modules with at least one human-duplicated gene (Table S2A, Figures 2D and S2C, Data S1). Of these, only the B-turquoise module, exhibiting low expression in early development and increases during cortical specification and the beginning of deep-layer formation (16 to 20 post-conception weeks), was enriched for human duplicated genes (n=76; *p*-value=4.17×10^-9^), in addition to autism-associated genes^68^ (n=61; *p*-value=1.86×10^-9^). Co-expressed genes within this module included several markers of GABAergic interneurons (e.g., *DLX1* and *DLX2*^69,70^) and deep-layer excitatory neurons (*SSTR2*^71^), with overrepresentation of GO terms related to neurogenesis, axonogenesis, and dendrite development (Table S2B). Based on these results, we next used transcriptomes from *ex-vivo*-induced neurons modeling human prenatal prefrontal cortex from pluripotency to upper layer formation (CORTECON^64^) to identify 21 WGCNA modules (Table S2C, Figures 2F and S2D, Data S1). Considering co-expression patterns within gene families, paralogs largely belong to different CORTECON modules with only six duplicate gene families in complete concordance (all paralogs in the same module) (Table S2D, Figure 2G). This demonstrates that our approach largely distinguishes transcriptional profiles between similar paralogs, and that expression diverges at relatively short evolutionary time scales (<6 million years), as we have shown for a smaller set of genes^72^.

Five CORTECON modules were significantly enriched for paralogs from human-specific gene families (*p*-values<0.05; Figures 2F and S2D)—highlighting neural functions related to stem cell population maintenance, intracellular receptor signaling, microtubule-based movement, and regulation of neuron projections (GO enrichments in Table S2E)—and including *SRGAP2C*, a human-specific gene known to interact with F-actin to produce membrane protrusions required for neuronal migration and synaptogenesis^73^ (C-blue module). Three modules were also enriched for autism-associated genes^68^ (*q*-value<0.05; Figure 2F), including the C-yellow module (n=20 candidate genes) associated with axon guidance and synaptogenesis (Table S2E). Duplicated genes in this module include *KANSL1*, the causal gene in Koolen-de Vries syndrome^74^, and others within autism-associated genomic hotspots (e.g., *CASTOR2*, *PMS2P6* and *STAG3L1* at the Williams-Beuren syndrome region (OMIM:609757)) making them compelling candidates in contributing to neurological features in children carrying CNVs at these loci (Figure S2E).

Collectively, our analysis identifies co-expression modules enriched for human-duplicated genes, such as the B-turquoise and C-blue modules, which both relate to regulation of neuron projection development. Additionally, we provide a complete list of duplicate paralog module assignments using data from post-mortem brain tissue and *in vitro* induced neurons to provide clues of their putative functions in brain development. Paralogs co-expressed in modules enriched for genes with known links to neurodevelopment and autism represent top candidates for follow-up experiments.

### Modeling functions of duplicated genes in brain development

The next step in understanding the role of human-duplicated genes in brain development is to test their functions in model systems. Our combined analysis highlights 148 gene families with at least one CN-constrained and brain-expressed human-duplicated paralog, in addition to 30 paralogs not assigned to a family (Table S1A, Figure 3). Of these, we found 106 with a homologous gene(s) in either mouse or zebrafish (Table S3A). Using matched brain-expression data from these species corresponding to human developmental stages^65,75,76^ (Figure S3, as previously described^77,78^) narrowed in on 76 and 41 single-copy orthologs expressed during neurodevelopment in mice and zebrafish (Table S3B), respectively, enabling functional studies. This leaves 40% of the human-duplicated families with no obvious mouse/zebrafish ortholog, including fusion genes, primate-specific genes (e.g., *TBC1D3* paralogs^25,79^), or those associated with great ape ancestral “core” duplicons^80^ (e.g., *NBPF* and *NPIP*). Alternative models are required, such as *in vivo* primate or cell culture organoids, to test the functions of these genes.

**Figure 3.**
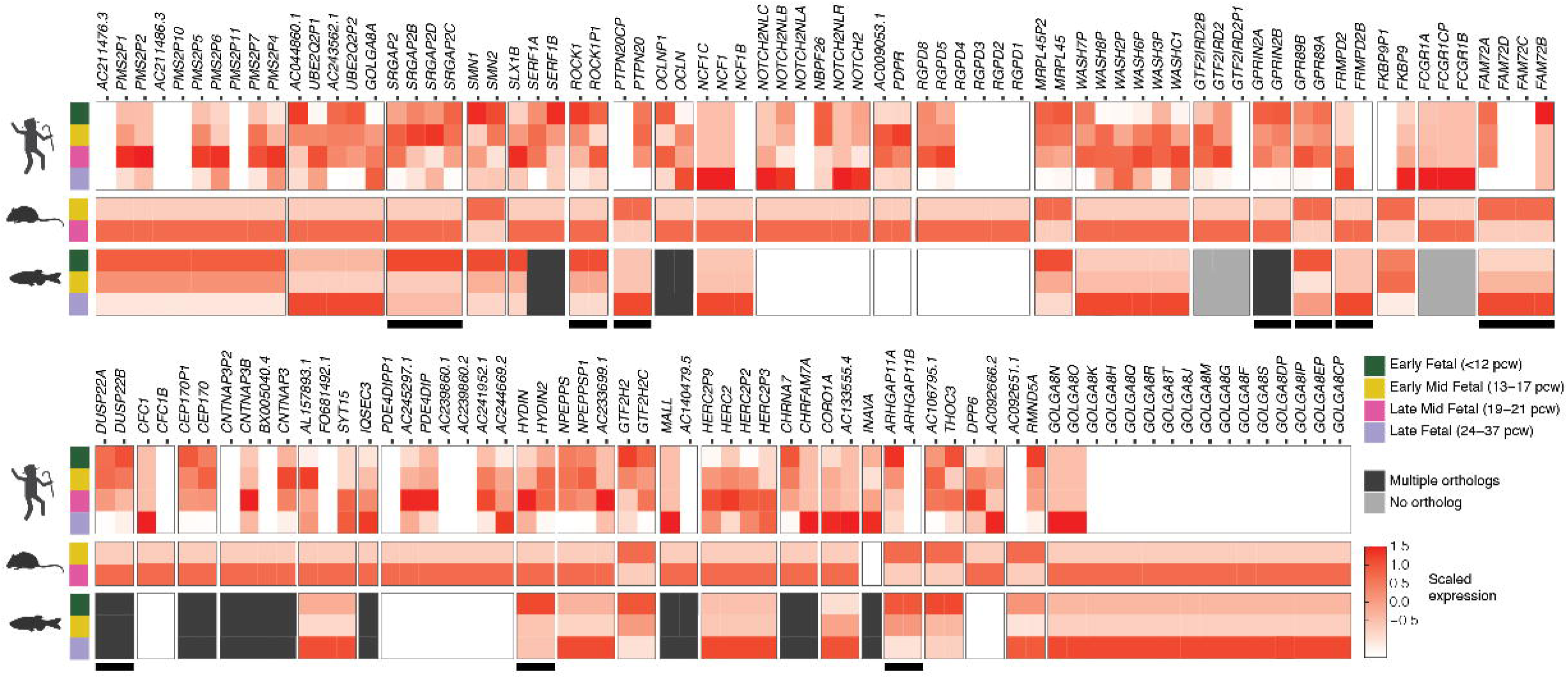
Modeling functions of duplicated genes in brain development. Scaled TPMs from the human BrainSpan dataset, and pseudo-bulk single-cell transcriptomes from whole-brain dissected samples of mouse and zebrafish. Gene families pictured represent a subset of CN-constrained and brain-expressed human-duplicated gene families with those highlighted with black bars prioritized for additional characterization. Also see Table S3 and Figure S3.

### Application of the resource: Characterizing candidate duplicated genes

#### Genetic variation of candidate genes important in neurodevelopment

As a proof of concept, we selected 13 priority human-specific duplicated (pHSD) gene families representing 30 paralogs from our model gene list (Table S4A, Figure 3). Since none of the paralogs fully reside within short-read-accessible genomic regions due to their high identity, we used published draft assemblies (112 total haplotypes)^81–84^ from the Human Pangenome Reference Consortium (HPRC) and Human Genome Structural Variation Consortium (HGSVC; Figure S4A) and performed capture high-fidelity (cHiFi) sequencing of 144 unrelated individuals of diverse ancestries^55,85^ (Table S4B–E, Figure S4B–E; see STAR Methods for details) to identify 46,754 variants (33,774 SNVs and 12,980 indels), or 12.7 variants/kbp, across targeted regions. Levels of variation within gene families were largely different between paralogs (Mann-Whitney U test, *p*≤0.05), with the exception of *FRMPD2* and *PTPN20* (Figure S4F). For instance, compared with the ancestral *SRGAP2* paralog, human-specific *SRGAP2B* exhibited the lowest and *SRGAP2C* the highest heterozygosity levels, in line with different mutation rates previously observed at each loci^18^.

Functional annotation^86^ identified 412 gene-impacting variants (Table S4F,G, Figure 4A), with eleven paralogs exhibiting no likely gene-disruptive (LGD) variants suggesting strong selective constraint. Virtually all paralogs had Ka/Ks lower than one, suggesting purifying selection, with seven ancestral and three derived paralogs exhibited Ka/Ks below the genome-wide average (∼0.25)^87^. The ancestral paralogs exhibited significantly lower Ka/Ks values than their derived paralogs (Wilcoxon signed-rank test, *p*-value=0.03) (Figure 4B), consistent with stronger purifying selection. More recent selection signatures incorporating polymorphic variation (pN/pS and the direction of selection [DoS]^88^) similarly indicated stronger purifying selection in the ancestral versus derived paralogs (Wilcoxon signed-rank test, *p*-value=0.023) (Figure 4C, Table S4H). While tests mostly agree, *NPY4R* shows discordant signatures, being highly conserved according to Ka/Ks but approaching zero in DoS, in line with an excess of observed LGD variants suggesting recent neutral evolution. Most paralogs within gene families have patterns expected under purifying selection, including *GPR89*, *CD8B*, *DUSP22*, *GPRIN2*, and *ARHGAP11* (also evident from a larger phylogenetic analysis of dN/dS using a maximum likelihood approach^89^; Table S4I).

**Figure 4.**
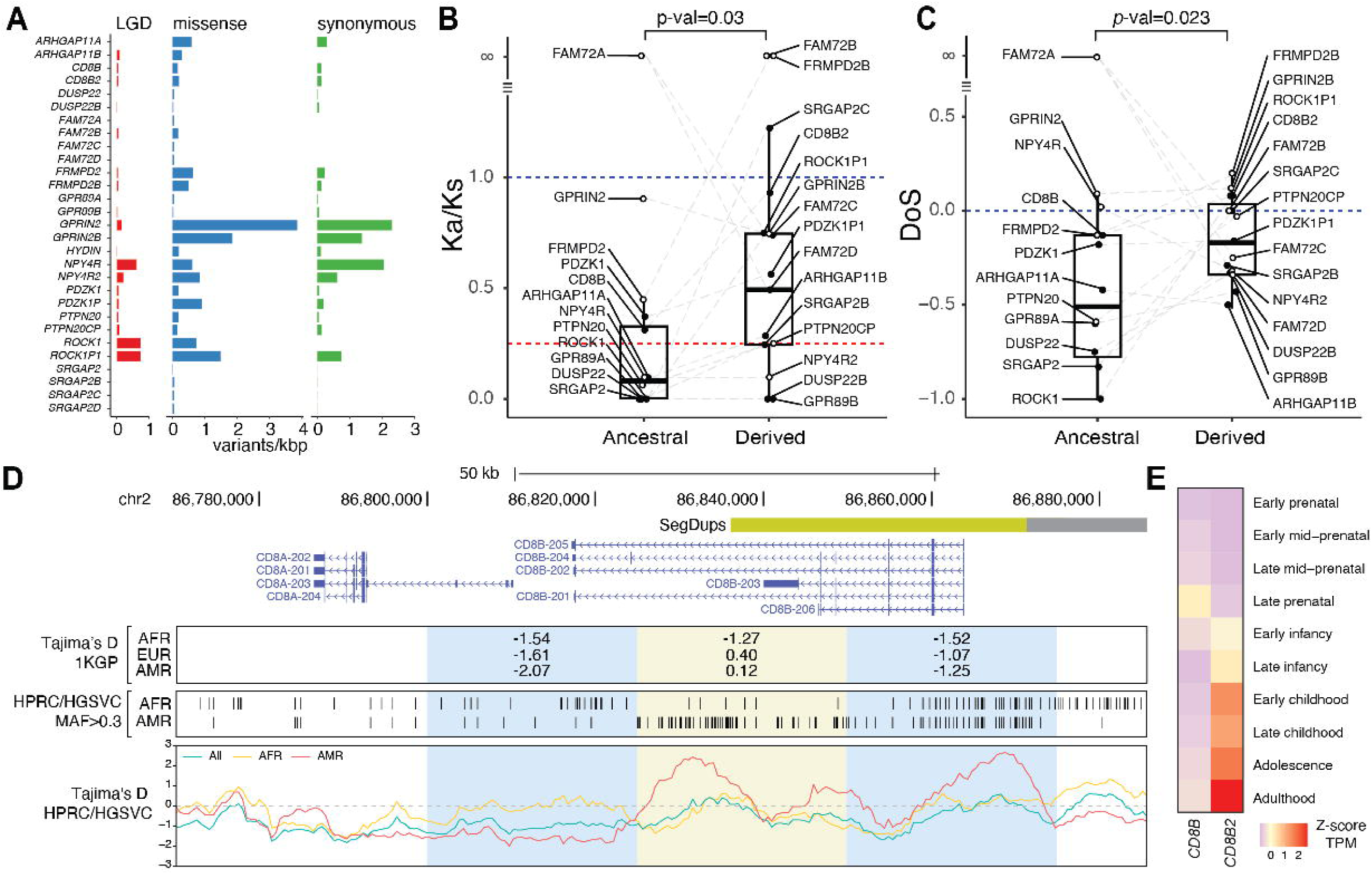
Genetic variation and signatures of selection of top candidate human-duplicated genes. (**A**) Number of likely gene-disruptive (LGD) (red), missense (blue), and synonymous (green) variants identified in pHSD genes. (**B**) Ka/Ks and (**C**) direction of selection (DoS) of pHSD genes with dashed lines indicating average genome-wide values between humans and chimpanzees (red) and neutrality (blue). Differences between matched ancestral and derived paralogs were tested with the Wilcoxon signed-rank test. Paralogs with infinite values or undetermined ancestral/derived state (hollow dots) were excluded from comparisons. (**D**) *CD8B* locus overview, including Tajima’s D values derived from 1KGP genome-wide SNVs (top panel). Biallelic SNVs from the Human Pangenome Reference Consortium (HPRC) and the Human Genome Structural Variation Consortium (HGSVC) assemblies are shown with with a minor allele frequency greater than 0.3 in individuals of African (AFR, n=27) and American (AMR, n=18) ancestry (middle panel) and used to calculate Tajima’s D values (bottom panel). (**E**) Scaled transcript per million (TPM) expression of *CD8B* and *CD8B2* in postmortem brain tissue from BrainSpan. Also see Table S4 and Figure S4.

Human-specific *SRGAP2C* has elevated Ka/Ks and pN/pS, together with low Tajima’s D in African individuals from the 1KGP genome-wide screen (−2.32; Figure 1E) suggesting positive selection, which we validated using high-confidence variants obtained from genome assemblies (−2.14; Figure S4G).

*GPR89* gene family paralogs exhibit low nucleotide diversity π and negative Tajima’s D values consistent with functional constraints (Figure S4H). *ROCK1* showed reduced π and more negative Tajima’s D compared to *ROCK1P1*, consistent with their divergent Ka/Ks values (Figure S4I). While Ka/Ks was not calculated for *FAM72* paralogs due to a lack of synonymous polymorphisms, Tajima’s D values similarly ranged from −2 to −1 indicating conservation of the gene family members (Figure S4J). Revisiting the 1KGP genome-wide signal of balancing selection in individuals of American and European ancestries centered on *CD8B* (Figures 1E and S1I), we find positive Tajima’s D in American (max 2.66, n=18) but not in African ancestries (max 0.62, n=27) (Figure 4D), evident as distinct haplotype clusters (Figure S4K).

The ancestral *CD8B* paralog, encoding CD8 Subunit Beta, is highly expressed in T cells where the protein dimerizes with itself or CD8A (alpha) to serve as a cell-surface glycoprotein mediating cell-cell interactions and immune response^90,91^. Considering all variants identified using assemblies and cHiFi sequencing, we observe an increase in intermediate-frequency variants, a signature of balancing selection, in *CD8B* among European and American ancestries, compared with those of African ancestry (Figure S4L, Kolmogorov-Smirnov, *p*-value=2.2×10^-16^), that differentiate the two main haplotypes. Two of the SNPs (rs56063487 and rs6547706) are *CD8B* splice eQTLs in whole blood from GTEx^92^ and significantly associated with increased CD8-protein levels on CD8+ T cells within a Sardinian cohort^93^.

We note that *CD8B2* paralog-specific variants do not overlap with the SNPs, providing confidence in these short-read-based genotype results. The haplotypes may, thus, play a role in the modulation of the adaptive immune response, a frequent target of balancing selection. The human-specific paralog *CD8B2* exhibits divergent expression in the human postnatal brain rather than in T cells^39^ (Figure 4E). These results provide an example of two paralogs with likely divergent functions and contrasting evolutionary pressures over a relatively short evolutionary time span (∼5.2 million years ago [mya]^14^). Combined, we demonstrate the efficacy of long-read data to uncover hidden signatures of natural selection.

#### Duplicated gene functions modeled using zebrafish

We performed a high-throughput functional screen in zebrafish^94–96^ of seven largely uncharacterized pHSD families expressed in both human and zebrafish brain^97,98^ (*GPR89*, *NPY4R*, *PTPN20*, *PDZK1*, *HYDIN*, *FRMPD2,* and *FAM72*; Table S3B, Figures 3 and S5A–C). Additionally, we tested two gene families (*SRGAP2* and *ARHGAP11*) previously studied in mammals^17,22–24,73,99–103^. While a whole-genome teleost duplication resulted in ∼20% of genes with multiple zebrafish paralogs that might confound functional analysis of human gene duplications^104^, the nine prioritized gene families tested here were selected in part because each had only one zebrafish ortholog (Table S3A). Ancestral gene functions were assessed using loss-of-function knockouts resulting in ∼70% ablation of alleles in G_0_ lines^105^ (termed crispants; Table S6 and STAR Methods) except for *arhgap11*, which is maternally deposited^106^, prompting us to use a morpholino that impedes translation. We also ‘humanized’ zebrafish models by introducing mRNAs encoding human-specific paralogs (Figure 5A) into wild-type embryos for all genes except *HYDIN2*, due to its large size (4,797 amino acids [aa]). This produced transient and ectopic presence of the transcript, detectable by RT-PCR at 3 dpf (Figure S5D). There were no significant morbidity differences in any tested models compared to controls (log-rank survival tests *p*-values>0.05, Table S5B).

**Figure 5.**
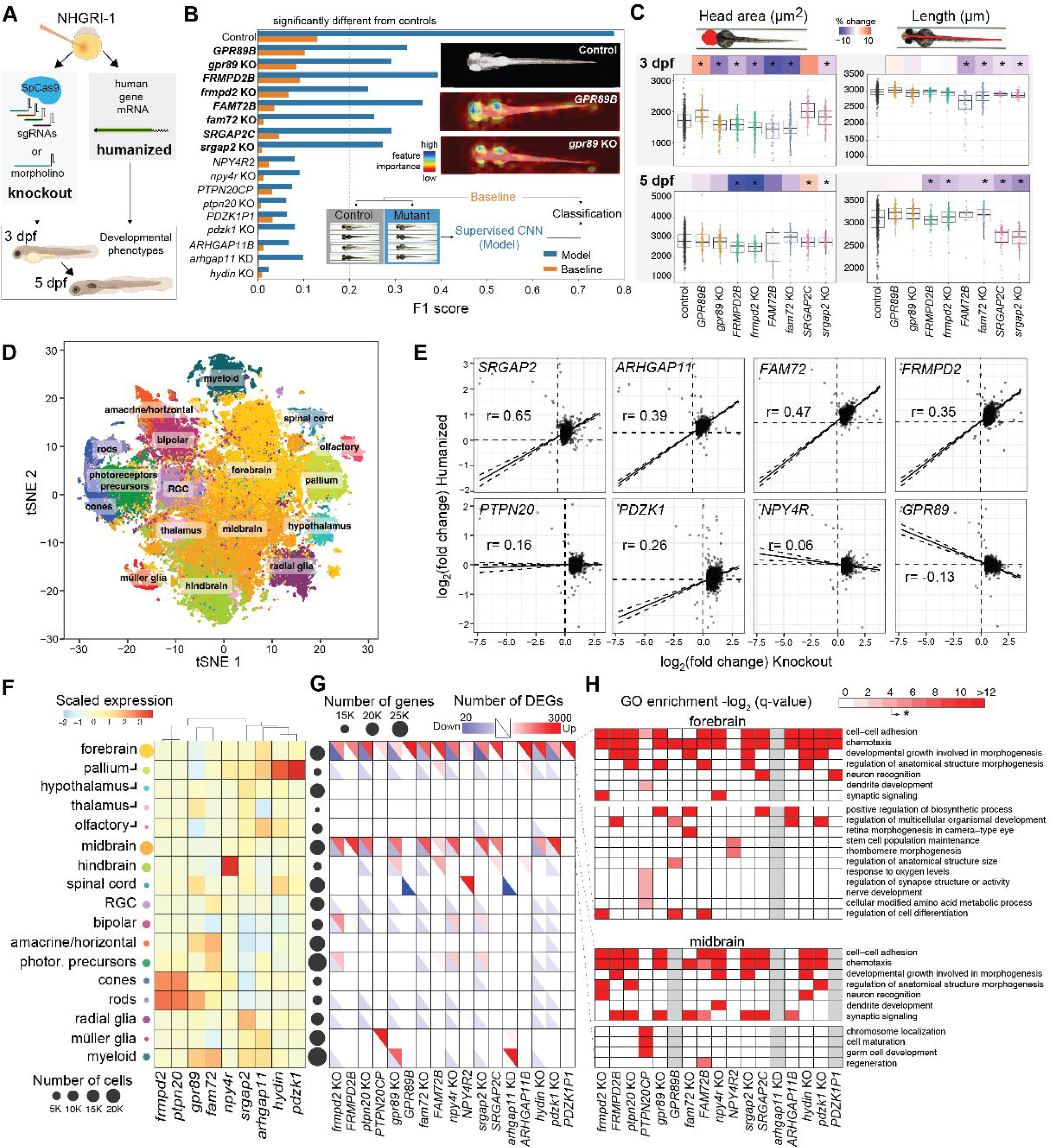
Duplicated gene functions modeled using zebrafish. **(A)** Functions of each pHSD gene were tested by generating knockout (KO, or morpholino) and ‘humanized’ models (injection of mRNA). **(B)** The F1 score, generated using a supervised convolutional neural network (CNN), is plotted indicating the effect size of morphological difference between models and controls, either using our batch-corrected images (blue bars) or original data (orange bars). Higher F1 score indicates greater difference. The bars for the control group indicate on average how distinct the controls are from all other groups. A threshold F1 score of 0.2 was used to define models being robustly classified as different from their control group. Pictured as a top inset are feature attribution plots with colors highlighting the region of the image used by the CNN to correctly classify and distinguish those genotypes from controls. **(C)** Measurements of selected pHSD gene families with heatmaps representing the percent change compared to the control group (asterisks indicating a Benjamini Hochberg-corrected *p*-value<0.05). **(D)** t-distributed stochastic neighbor embedding (tSNE) plot highlighting classified cell types from scRNA-seq data at 3 dpf. **(E)** Fold-change comparison between KO and humanized models for each pHSD across all genes (n= 29,945), versus their controls. Black lines represent the Pearson correlation line and the dotted lines the 95% confidence intervals. **(F)** Endogenous z-score scaled expression of each zebrafish ortholog across defined scRNA-seq cell types. Circle sizes scale with the overall number of cells included in that group. **(G)** Distribution of cell-type-specific differentially expressed genes (DEGs) for each pHSD model. Each square includes the downregulated genes in blue (lower diagonal) and upregulated genes in red (upper diagonal). Circles next to each cell type represent the number of expressed genes. **(H)** Gene ontology (GO) enrichment results for the top overrepresented terms in upregulated genes in forebrain and midbrain across pHSD models, with gray indicating genes with no DEGs. Significant q-value>0.05 indicated with asterisk on color legend. Also see Table S5 and Figure S5.

Significant morphological differences were first identified without predefining specific features *a priori* by imaging^107–109^ at 3 and 5 days post-fertilization (dpf) and using latent diffusion and convolutional neural networks (CNNs)^110^ (Table S5C, STAR Methods). This flagged knockout and humanized models of *SRGAP2*, *GPR89*, *FRMPD2*, and *FAM72* (F1 scores>0.2, Figure 5B, STAR Methods). Quantifying specific features using the same images (Table S5D,E) revealed concordant phenotypes for knockout and humanized models of *SRGAP2* (reduced length), *FRMPD2* (reduced head area), and *FAM72* (both reduced body length and head area) at 3 dpf (Figure 5C). Alternatively, *GPR89* models exhibited opposing effects, with head area for *gpr89* knockout larvae ∼10% reduced and *GPR89B* ‘humanized’ larvae ∼15% increased. This is also evident in the feature attribution plot indicating that the CNN distinguishes both *gpr89* knockout and *GPR89B* humanized larvae from controls primarily by focusing on the head (Figure 5B). At 5 dpf, the alterations in *FRMPD2* and *SRGAP2* models persisted while no longer observed for *FAM72* and *GPR89* (Figure 5C). Knockout models for *gpr89* and *frmpd2* also displayed evidence of developmental delay with subtle yet significant decreases in the head-trunk angle^111^.

To directly characterize impacts on brain development, we profiled 95,555 cells (an average of 3,822±3,227 per model; Figure S5E) using single-cell RNA-sequencing (scRNA-seq)^112,113^ from dissected heads of 3 dpf larvae (Figure 5D). Pseudo-bulk differential expression analysis of all cells revealed significant positive correlations for *SRGAP2C*, *FAM72B*, *ARHGAP11B*, *FRMPD2B*, and *PDZK1B* humanized larvae with respect to each knockout indicating loss-of-function effects (Figure 5E). *GPR89B* gene expression changes are negatively correlated with *gpr89* indicating gene dosage effects, while *PTPN20CP* and *NPY4R2* show low/no relationship between models. These results are in line with our morphometric findings for *SRGAP2*, *FRMPD*2, *FAM72*, and *GPR89* (Figure 5C), as well as from our separate study^114^ that verified the human SRGAP2C protein physically interacts with and antagonizes zebrafish Srgap2. The concordant phenotypes for gain- and loss-of-function for over half of the tested genes fits the gene-balance hypothesis^115,116^ and recent results in humans showing that large increases and decreases in gene expression via CNVs can impact certain complex traits in the same direction^117^.

Classifying 17 different neuronal, retinal, and glial cell types^76,112,118,119^ shows pHSD orthologs are broadly expressed, with a subset showing more narrow expression patterns (e.g., *hydin* and *pdzk1* in the pallium, *npy4r* in the hindbrain; Table S5F,G, Figure 5F). Pseudo-bulk analyses revealed gene dysregulation in most cell types (Figure 5G; DEGs available here^57^). GO enrichment of DEGs in the forebrain (the closest related structure to the human cerebral cortex^120^) and midbrain (the main visual processing center primarily consisting of the optic tectum), where we had the greatest power to detect differences due to the largest abundances of cells, suggests alterations in cell-cell adhesion, chemotaxis, and altered synaptic signaling (Figure 5H, Table S5H). Several humanized models exhibited unique effects in DEGs, such as in the midbrain of *GPR89B* models and regulation of anatomical structure size, Müller glia in *PTPN20CP*, and the spinal cord in humanized *NPY4R2*. Myeloid cells were also uniquely impacted in *gpr89* and *arhgap11* knockout larvae. Combined, these results indicate that all tested pHSD models impact the developing zebrafish brain, suggesting that they may also play important roles in human brain evolution.

#### Human-specific genes impacting neurodevelopment

##### *GPR89B* and brain size

Opposite phenotypes were observed for *gpr89* knockout and humanized *GPR89B* zebrafish suggesting gene dosage effects. Considering both *GPR89* human paralogs are impacted by deletions and duplications at the chromosome 1q21.1 genomic hotspot associated with microcephaly and macrocephaly in children with neurocognitive disabilities, respectively^59^, we sought to characterize mechanisms underlying larval head-size phenotypes in more detail. Generating a stable *gpr89* mutant line (STAR Methods) showed that heterozygous and homozygous knockouts exhibited reduced head size at 3 dpf, verifying results in crispants (Figures 5C and 6A). Further, we observed significantly smaller and larger forebrains in crispant G_0_ knockout and humanized zebrafish larvae, respectively, using a neuronal reporter line^121^. Sub-clustering cells from the forebrain, we observed endogenous expression of *gpr89* in telencephalon and inner diencephalon (Figure 6B). DEGs with inverse effects between *GPR89* knockout and humanized models in the telencephalon, a brain structure anatomically equivalent to the mammalian forebrain with roles in higher cognitive functions such as social behavior and associative learning^122,123^, were enriched in negative regulation of the DNA replication and cell cycle (Figure 6C, Table S6AB). Several genes functioning at the G2/M checkpoint were downregulated in the humanized *GPR89B* and upregulated in the knockout *gpr89* pointing to differences in cell proliferation. We estimated the identity of forebrain cells and found humanized *GPR89B* cells more likely to classify as neural progenitors while *gpr89* knockouts more likely to be differentiated neurons (Figure 6D).

**Figure 6.**
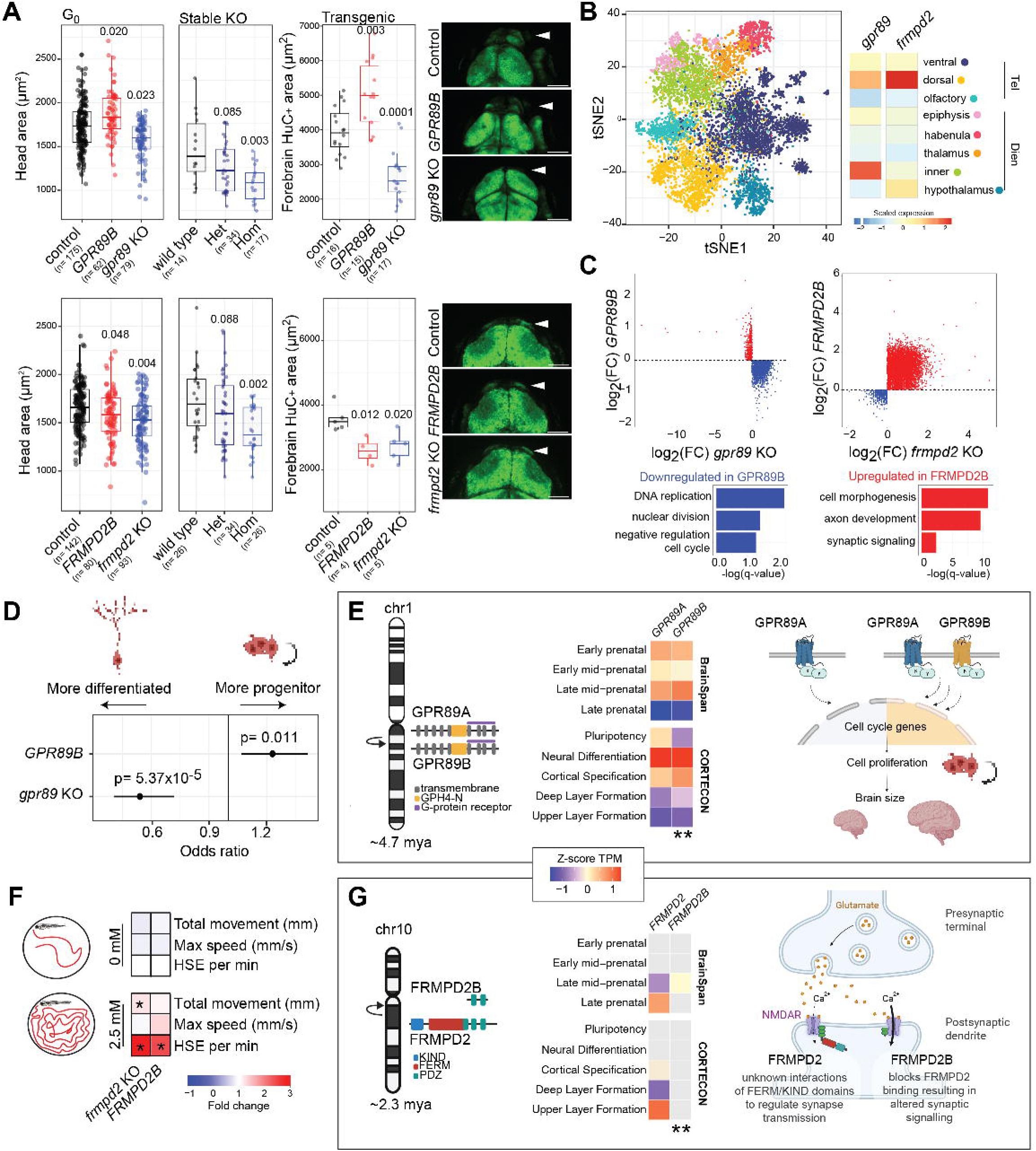
Neurodevelopmental impact of *GPR89* and *FRMPD2*. **(A)** Head and brain area assessments at 3 dpf for G_0_ crispants and stable knockout lines. *p*-values are indicated above box plots versus controls using an ANCOVA with a rank-transformation (humanized and crispant models) and Wilcoxon signed-rank tests (stable knockout lines). Representative images of each model in the neuronal transgenic line are included with scale bars representing 100 µm. **(B)** t-distributed stochastic neighbor embedding (tSNE) plot showing the identified subregions classified from the forebrain (n=10,040 cells) and relative scaled endogenous expression across cell types. **(C)** Log_2_ fold change (FC) of gene expression versus controls in cells from the telencephalon between knockout and humanized models. Red and blue colors correspond to DEGs discordant (*GPR89*) or concordant (*FRMPD2*) between the knockout and humanized models and their top representative gene ontology (GO) enrichment. **(D)** Forest plot with the results from the logistic regression for presence of progenitor versus differentiated states in forebrain cells. **(E)** Diagrams of the duplication event of *GPR89* with different expression patterns (**Wilcoxon signed-rank test, *p*-value<0.005). A model of GPR89B gain-of-function in neuronal proliferation amplification is depicted on the right. **(F)** Behavioral results from 1 h motion-tracking evaluations in 4 dpf larvae exposed (2.5 mM) or not (0 mM) to pentylenetetrazol (PTZ) with high-speed events (HSE) defined as movement ≥28 mm/s. Colors represent the FC relative to the control group and the asterisk indicates a significant Dunn’s test (*p*<0.05 Benjamini-Hochberg-adjusted). **(G)** Diagram of the duplication event of *FRMPD2* (see also **E**), with a model of FRMPD2B antagonistic functions resulting in altered synaptic signaling depicted on the right. Also see Table S6.

*GPR89* (G-protein receptor 89 or *GPHR*, Golgi PH regulator) encodes highly conserved transmembrane proteins that participate in intracellular pH regulation in the Golgi apparatus^124^. Loss of function in *Drosophila* leads to global growth deficiencies due to defects in the secretory pathway^125^. In humans, a complete duplication of ancestral *GPR89A* ∼4.7 mya produced a derived *GPR89B*^14^ (Figure 6E). The two paralogs maintain identical protein similarity but differential and overlapping expression patterns in human brain development, with *GPR89A* evident in pluripotency (C-turquoise module), and *GPR89B* expression turning on slightly later during neural differentiation (C-red module; Figures 2F and 6E). Both genes show evidence of purifying selection (Figure S4), with *GPR89A* exhibiting extreme negative Tajima’s D values in individuals of African and American ancestries from the 1KGP cohort (<5^th^ percentile; Figure 1E). These results suggest that *GPR89* paralogs function in early brain development, with delayed expression of *GPR89B* possibly extending expansion of progenitor cells, a feature observed in human cerebral organoids compared with those of other apes^126,127^ (Figure 6E). Together with the increase in forebrain size of “humanized” zebrafish, we propose a role for *GPR89B* in contributing to the human-lineage expansion of the neocortex.

##### *FRMPD2B* and synaptic signaling

While opposing traits were observed in *GPR89* models, similar phenotypes suggest that the human FRMPD2B acts as a dominant negative to the endogenous Frmpd2. We generated a stable *frmpd2* mutant line (STAR Methods) and observed reduced head size in homozygous knockout larvae validating crispant features (Figures 5C and 6A). Additionally, both the crispant knockout *frmpd2* and humanized *FRMPD2B* larvae exhibit smaller forebrains. Shared upregulated DEGs between *FRMPD2* models function in cell/axon morphogenesis and growth as well as synaptic signaling in telencephalic cells (Figure 6C, Table S6C,D). To better characterize impacts on synaptic signaling, we treated with a low dose of the GABA-antagonizing drug pentylenetetrazol (PTZ) producing a significant increase in high-speed events^128^, indicative of seizures in larvae, in both humanized and knockout *FRMPD2* larvae (4 dpf) versus controls (Figure 6F). These results suggest that Frmpd2 loss of function, through *frmpd2* knockout or antagonism via FRMPD2B, disrupts synapse transmission and amplifies induced seizures, in line with the known interactions of FRMPD2 with glutamate receptors^129^.

*FRMPD2* (FERM and PDZ domain containing 2) encodes a scaffold protein that participates in cell-cell junction and polarization^130^ localized at photoreceptor synapses^131^ and the postsynaptic membrane in hippocampal neurons in mice^129^. A partial duplication of the ancestral *FRMPD2* created the 5’-truncated *FRMPD2B* paralog ∼2.3 mya^14^. *FRMPD2B* encodes 320 aa of the C-terminus, versus 1,284 aa for the ancestral, maintaining two of three PDZ domains involved in protein binding^132^ but lacking the KIND and FERM domains (Figure 6G). Ancestral *FRMPD2* expresses in the human prenatal cortex during upper layer formation, while *FRMPD2B* is evident postnatally^65^. The paralogs also show divergent evolutionary signatures, with *FRMPD2* strongly conserved and *FRMPD2B* exhibiting possible positive selection (Figure 4B,C). Combined, we propose a model in which truncated human-specific FRMPD2B counteracts the function of full-length FRMPD2 leading to altered synaptic features in humans, possibly through interactions of its PDZ2 domain with GluN2A of NMDA receptors at the postsynaptic terminal^129^. Its postnatal expression would avoid the detrimental effects of inhibiting FRMPD2 during early fetal development (i.e., microcephaly). We note that recurrent deletions and duplications in chromosome 10q11.21q11.23 impact both paralogs in children with intellectual disability, autism, and epilepsy^133^. Ultimately, *FRMPD2B* could plausibly contribute to the upregulation of glutamate signaling and increased synaptic plasticity observed in human brains compared with other primates that is fundamental to learning and memory^134^.

## Discussion

Our results provide the scientific community with a prioritized and comprehensive set of hundreds of genes to perform functional analyses with the goal to identify drivers of human brain evolution (213 families and 1,002 total paralogs). Compared to a previous assessment of human-specific duplicated genes^14^, this represents an approximately fivefold increase, in part because we also included human-expanded gene families and genes with as little as one duplicated exon. These numbers are likely an underestimate, as we excluded 193 high-copy gene families (famCN>10), as well as families that have undergone independent gene expansions or incomplete lineage sorting with other great apes. One compelling example is *FOXO3*, encoding the transcription factor forkhead box O-3, implicated in human longevity^135^, with all three paralogs CN-constrained and brain expressed. Since this gene also exists as duplicated in other great apes at similar CN, we excluded it from our list of human gene expansions. This is, in part, because there is still uncertainty regarding which paralog(s) are human specific due to secondary structural rearrangements that hamper genomic alignments^68,136^. Moving forward, the availability of nonhuman primate T2T genomes will improve orthology and synteny comparisons between species^137–139^. As a resource for the community, we have made available the results of our genome-wide analyses across the complete 1,793 SD98 genes (Table S1A).

Collectively, 148 gene families (362 paralogs, 108 annotated as non-syntenic with the chimpanzee reference) represent top candidates for contributing to human-unique neural features. In this study, we chose zebrafish to demonstrate the efficacy of our gene list. Despite notable differences with humans, such as the absence of a neocortex^140^, conservation in major brain features make zebrafish well suited to characterize gene functions in neurological traits, including cranial malformations^141^, neuronal imbalances^142^, and synaptogenesis^143^. Coupled with CRISPR mutagenesis^94,95^, zebrafish have been used as a higher-throughput model for human conditions such as epilepsy^128^, schizophrenia^144^, and autism^78^.

From our analysis, knockout and humanized models of four genes (*GPR89*, *FRMPD2*, *FAM72*, and *SRGAP2*) resulted in altered morphological features, primarily to head size (often used as a proxy for brain size), and all models exhibited molecular differences in single-cell transcriptomic data (Figure 5G). Two duplicate gene families, *SRGAP2* and *ARHGAP11*, have been extensively studied in diverse model systems (reviewed recently^9^). Our zebrafish model of *SRGAP2*, encoding SLIT-ROBO Rho GTPase-activating protein 2, were consistent with published findings in mouse where the 3’-truncated human-specific SRGAP2C inhibits the function of the endogenous full-length Srgap2^17^. Further, the shared upregulated genes identified in the forebrains of *SRGAP2* mutant larvae point to alterations in axonogenesis and cell migration (Table S5I), matching studies in mice^11,17,18,73,100,145,146^. Alternatively, *ARHGAP11B*, encoding Rho GTPase Activating Protein 11, implicated in the expansion of the neocortex through increased neurogenesis^22,24^, exhibited no detectable changes in head/brain size when introduced in zebrafish embryos. Upregulated DEGs were only detected in the forebrains of *ARHGAP11B*-injected mutants and were enriched in cellular biosynthetic processes (mRNA splicing and translation; Table S5J). Given that ARHGAP11B impacts the abundance of basal progenitors, a cell type unique to the mammalian neocortex^147^, zebrafish may not be suitable to characterize human-specific functions of this gene. This was also evident in *ARHGAP11B*-humanized zebrafish that exhibited similar molecular changes to *arhgap11* morphants (Figure 5E) suggesting antagonism by the human paralog, which is counter to previous studies^148^ and possibly a spurious result due to the ectopic human mRNA expression.

Beyond modeling gene functions, our study also highlighted the considerable amount of genetic variation hiding within SD regions. Even with a complete T2T-CHM13, only 10% of SD98 regions are “accessible” to short reads^37^ resulting in <10% sensitivity to detect variants and a depletion of GWAS hits (STAR Methods). Analysis of existing assemblies (HPRC and HGSVC) and cHiFi sequencing uncovered some of this hidden variation within 13 pHSD gene families. We note that for some highly identical duplicated genes (*CFC1*), cHiFi reads (∼3 kbp) were still too short to accurately map to respective paralogs (data not included). Nevertheless, long reads revealed that most pHSD paralogs exhibit evolutionary constraints and provided support for balancing selection of *CD8B*, not previously identified in published genome-wide screens^149,150^. Historically, signatures of balancing selection, which include an excess of mid-frequency alleles^151^, have been difficult to detect within SDs due to assembly errors^37^. In these cases, paralog-specific variants are mistaken for SNPs when reads from both paralogs map to a single collapsed locus resulting in false mid-frequency alleles. Scientific consortia like *All of Us* are generating long-read datasets at scale^152^, ushering in a new era where genomic associations and evolutionary selection may finally be uncovered within human duplications to identify novel drivers of human traits and disease.

Similarly, genome sequencing of patients and their families has discovered hundreds of compelling neuropsychiatric disease candidate genes impacted by rare and *de novo* variants, but the genetic risk underlying conditions such as autism is still not completely elucidated^153^. SD genes may represent a hidden contributor to disease etiology. Our analysis identified 231 SD98 genes (110 human duplicate paralogs) co-expressed in modules enriched for autism genes (Table S2), including several within disease-associated genomic hotspots. Distinct SD mutational mechanisms, including ∼60% higher mutation rate compared to unique regions^154^ and interlocus gene conversion that can occur between paralogs^155,156^, make duplicated genes particularly compelling to screen for *de novo* mutations contributing to idiopathic conditions. For example, nonfunctional paralogs with truncating mutations can “overwrite” conserved functional paralogs leading to detrimental consequences, as is the case of *SMN1* and *SMN2* in spinal muscular atrophy^40^. Human-duplicated gene families include ancestral paralogs *CORO1A*, *TLK2*, and *EIF4E*, with significant genetic associations with autism^68^. We propose that interlocus gene conversion between their likely nonfunctional duplicate counterparts is an understudied contributor to neurodevelopmental conditions in humans. Our comprehensive list of gene families will enable future work to progress in this research area.

Our study focuses on brain development, but primates exhibit other prominent differences across musculoskeletal and craniofacial features that have diverged early in human evolution^4^. Since such traits are largely universal across modern humans, our list of CN-constrained genes represent top candidates but re-analysis of transcriptomes from non-brain cells/tissues is required. Meanwhile, duplicate genes, such as those encoding defensins^157–160^, mucins^161,162^, and amylases^42–44^, can also play a role in metabolism and immune response that exhibit population diversification due to the vast variability in diet, environment, and exposures to pathogens across modern humans^28^. Our use of a single human T2T-CHM13 haplotype of largely European ancestry^35^ could miss some CN-polymorphic genes. As additional T2T genomes are released^29^, it will be important to continue curating this list of duplications. Nevertheless, genes CN stratified by human ancestry can be identified using metrics such as V ^163^, as has been highlighted in other studies (reviewed here^9^ and most recently^164^). Facilitating such analyses for our gene set, we provide a publicly available resource to query parCN median estimates across individuals from 1KGP for our complete set of SD98 paralogs (https://dcsoto.shinyapps.io/shinycn).

### Limitations of the study

A notable limitation of our study is its reliance on existing gene annotations, which we used to group human duplicate paralogs into larger multigene families based on shared annotated sequences in SD98 regions. Due to the complexities of SDs, which can result in gene fusions and altered gene structures, some genes were left unassigned to a family (n=114 singletons from SD98 genes). Other noncoding transcripts and lncRNAs were excluded altogether, including a human-specific paralog of *IQSEC3*, a gene implicated in GABAergic synapse maintenance^165^. Additionally, the functional consequences of variants identified in 656 unprocessed pseudogenes are difficult to interpret. Improvements are on the horizon, with ongoing work with long-read transcriptomes that will continue to refine annotations^166^ and advancements in protein-prediction^167^ and proteomic approaches^168^ that will confirm whether or not these genes encode proteins. Similarly, single-cell transcriptomes typically focus on 3′-ends of transcripts, limiting specificity of human paralogs. Generation of single-cell long-read datasets^169^ will enable more refined assessments of duplicate genes to discern differences in expression across cell types in the brain and other tissues. Further, for this study we have focused our analysis on gene duplications, but other complex structural variants have high propensity in altering functions and/or regulation of genes^28^, with human-specific deletions^170,171^ and inversions^172^ the focus of recent studies. Finally, we highlight that our zebrafish studies employ transient, ectopic expression of human paralogs to “humanize” larvae and characterize phenotypes, which limited our analysis to early developmental traits (>4 dpf in zebrafish^141^), approximately equivalent to human mid-to late-fetal stages in brain development (Figure S3C), and could result in spurious phenotypes. Moving forward, generating stable transgenic zebrafish and mammalian models that better match endogenous cell/tissue expression of human paralogs will enable more precise delineation of gene functions.

In summary, we interrogated challenging regions of the genome by taking advantage of long-read sequencing in tandem with the new T2T-CHM13 reference genome and demonstrated a method using zebrafish to explore the functions of human-duplicated genes. Among our list of hundreds, we propose duplicate gene paralogs potentially contributing to unique features of the human brain, specifically featuring two: *GPR89B*, with a possible role in expansion of the neocortex, and *FRMPD2B*, with implications in altered synaptic signaling. In the future, additional genetic analyses across modern and archaic humans and experiments utilizing diverse model systems will reveal hidden roles of duplicated genes in human traits and disease.

## Resource Availability

### Lead contact

Further information and requests for resources should be directed to and will be fulfilled by lead contact Megan Y. Dennis.

### Materials availability

- Plasmids generated in this study have been deposited to Addgene.
- Zebrafish lines generated in this study *frmpd2^tupΔ^*^5^ and *gpr89^tupΔ^*^8^ are available from the lead contact upon request.

### Data and code availability

- All raw sequencing data generated from this study, including cHiFi sequencing and scRNA-seq from zebrafish, have been deposited to ENA and NCBI (PRJEB82358). Accession numbers of data used from published sources can also be found in the Key Resources Table.
- All original code and processed data associated with this study is publicly available through https://github.com/mydennislab/HSD_brain_evolution and https://github.com/Ricardo-scb/ZebraFish-Diffusion-Model/ and has been deposited at Zenodo (DOIs 10.5281/zenodo.15486469 and 10.5281/zenodo.15485460).
- Any additional information required to reanalyze the data reported in this paper is available from the lead contact upon request.

## Acknowledgements

We would like to thank the 1KGP, HGDP, T2T, SSC, gnomAD, HPRC, and HGSVC for access to their data and biospecimens. Thank you to Dr. Alfie Gleeson and Cassandra Olivas for support with evolutionary and transcriptome analyses, respectively, as well as Tonia Brown for copyediting. The Cell Biology and Human Anatomy Department at UC Davis provided significant imaging resources. This work was supported, in part, by U.S. National Institutes of Health (NIH) grants from the Office of the Director and National Institute of Mental Health (DP2MH119424 and R01MH132818 to M.Y.D., DP2MH129987 to G.Q.), National Institute of Neurological Disorders and Stroke grant (R01NS109176 to S.S.), National Institute of Child Health and Human Development (P50HD103526 to G.Q.), as well as National Science Foundation CAREER awards (2145885 to M.Y.D., 1846559 to G.Q.). Support for A.M.A. came from UCL Wellcome Trust ISSF3 award (204841/Z/16/Z) and Biotechnology and Biological Sciences Research Council, BBSRC (BB/WW007703/1). NIH training grants through the National Institute of General Medical Sciences supported A.S. (T32GM007377), N.K.H. (T32GM153586), and N.A.F.M. (R25GM116690). Some figures were made with Biorender.

## Author Contributions

D.C.S., J.M.U-S. and M.Y.D. conceived the project; D.C.S., J.M.U-S., G.K., A.S., and M.Y.D. generated sequencing data and analyzed datasets; D.C.S., A.S., A.M.A., and M.Y.D. performed genetic variation and population genetic analysis of human datasets; D.C.S. and Z.J. performed WGCNA analysis; R.V. and G.Q. performed supervised classification of zebrafish larvae; J.M.U-S, N.K.H., K.H., G.L. N.A.F.M., C.I., A.B., S.S., and M.Y.D. contributed to genotyping and phenotyping animal models associated with this study; T.N.T. generated the parCN dataset for the SSC cohort; E.D.G. provided human DNA samples; D.C.S., J.M.U-S., and M.Y.D. drafted the manuscript. All authors read and approved the manuscript.

## STAR Methods

### Experimental model and study participant details

#### Study participants

Sequenced genomes from human individuals were included from publicly available resources (1000 Genomes Project [1KGP] ^55^, Human Genome Diversity Project [HGDP] ^173^, Simons Genome Diversity Project [SGDP] ^45^, Genome in a Bottle ^174^, Human Pangenome Reference Consortium [HPRC)]^29^, and Human Genome Structural Variation Consortium [HGSVC] ^84^) or through controlled access (Simons Simplex Collection [SSC] ^60,61^) with known ancestries. Sex was not considered, since we focused on genetic variation across autosomes. This study was reviewed by the Institutional Review Board of the University of California, Davis and deemed minimal risk and human subjects exempt, with all participants de-identified.

#### Zebrafish procedures

Wild-type NHGRI-1^175^, Tg[HuC-GFP] ^121^, and mutant (generated from this study) adult zebrafish were maintained in a modular system (Aquaneering, San Diego, CA) distributed in tanks at a maximum density of 10 adults per L and all fish were kept in temperature (28±0.5°C) and light (10h dark / 14h light cycle) controlled environment following standard protocols ^176^ with flowing water filtered via UV (Aquaneering, San Diego, CA). As described previously ^105,109^, all animals were monitored twice daily for health evaluations and feeding that included brine shrimp (Artemia Brine Shrimp 90% hatch, Aquaneering, San Diego, CA) and flakes (Select Diet, Aquaneering, San Diego, CA). To obtain embryos for the different experiments, NHGRI-1 males and females of at least three months of age were randomly selected and placed in 1L breeding tanks in a 1:1 ratio and eggs from at least five crosses collected and kept in Petri dishes with E3 media (0.03% Instant Ocean salt in deionized water) in an incubator at 28°C at a density of less than 100 embryos per dish until used. Embryos from all breeding crosses were pooled together and for all experiments embryos were collected at random. CRISPR-generated *frmpd2^tupΔ^*^5^ and *gpr89^tupΔ^*^8^ stable mutant F_1_ zebrafish carrying heterozygous alleles were outcrossed twice to wild-type NHGR1-1 adults to remove potential off-target edits and then incrossed to generate batches of wild-type, heterozygous, and homozygous larval siblings for phenotypic assessments, following standard protocols^177^. All phenotypic screening was performed in zebrafish early larvae (up to 5 dpf). At this stage, animals are sexually indifferent. Sex determination in zebrafish is not well understood as it occurs in the absence of sex chromosome but it is generally accepted that sex determination establishes during the late larval stage of development (between approximately 15–25 dpf) ^178^. All animal use was approved by the Institutional Animal Care and Use Committee from the Office of Animal Welfare Assurance, University of California, Davis.

### Method details

#### Identification of SD98 genes

Duplicated regions were extracted from previously annotated SDs ^36^ using T2T-CHM13 (v1.0) coordinates and subsequently merged using BEDTools merge ^179^. SD98 regions were defined as an SD with ≥98% sequence identity to another locus in the T2T-CHM13 genome using the fractMatch parameter. Gene coordinates were obtained from T2T-CHM13 (v1.0) CAT/Liftoff annotations (v4) ^35^. SD98 genes were defined as gene annotations that contain at least one exon fully contained within an SD98 region, calculated with BEDTools intersect using -f 1 parameter ^179^. Overall numbers of distinct gene features overlapping SD98 were counted using the gene ID unique identifiers. We noticed that, in a few cases, two transcript isoforms of the same gene were assigned to different gene IDs. To identify these redundant transcripts, we self-intersected SD98 transcripts, selected those with different gene ID that also shared >90% positional overlap, and performed manual curation of the obtained gene list, removing redundant and read-through fusion transcripts. Gene coordinates were lifted over to the T2T-CHM13v2.0 assembly using UCSC liftOver tool ^180^ and the chain file for v1.0 to v1.1 conversion, obtained from the T2T-CHM13 reference GitHub repository (https://github.com/marbl/CHM13).

#### Gene family clustering

SD98 genes were grouped into gene families based on shared exons (Table S1C). Starting from T2T-CHM13 (v1.0) annotations, DNA sequences of all SD98 regions were extracted using BEDTools getfasta and mapped back to the reference genome using minimap2 (v2.17) ^181^ with the following parameters: -c - -end-bonus 5 --eqx -N 50 -p 0.5 -t 64. For each SD98 exon, the BEDTools intersect with -f 0.99 parameter was used to select mappings covering >99% of the exon sequence, removing self-mappings. This list was refined using the previously published ^36^ whole-genome shotgun sequence detection (WSSD) ^10^ CNs (famCN) of humans from the SGDP (n=269), which provides estimates of the overall CN of a gene family using read depth of multi-mapping reads with nonoverlapping sliding-windows. After comparing the median famCN values of SD98 genes with shared exons, groupings where the mean absolute deviation of the CN was less than one were selected. The list was filtered to focus on gene families containing at least one protein-coding or unprocessed pseudogene. SD98 genes associated with other gene features, including lncRNAs and processed pseudogenes, were also assigned a gene family ID. On the other hand, if a gene was not associated with any other gene feature, they were classified as “unassigned” or “singletons”. SD98 gene families were intersected with previously published DupMasker annotations using BEDTools intersect, which indicate ancestral evolutionary units of duplication ^36^.

#### Human-duplicated gene families

Human-specific and -expanded gene families were identified using CN comparisons between humans and nonhuman great apes with previously published WSSD ^10^ (famCN) CNs from humans (SGDP n=269) and four nonhuman great apes, including one representative of chimpanzee (Clint), bonobo (Mhudiblu), gorilla (Kamilah), and orangutan (Susie) ^36^, mapped to T2T-CHM13 (v1.0). The median famCN per SD98 gene was calculated using a custom Python script. For each SD98 gene, putative gene family duplications and expansion were predicted, excluding genes with median famCN>10 across humans from this analysis. Genes were considered expanded if the median famCN across humans was greater than the maximum famCN across great apes. Human duplications and expansions were distinguished based on whether the maximum famCN value across great apes was less than 2.5 (non-duplicated in great apes) or greater than 2.5 (duplicated in great apes), respectively. Non-syntenic paralogs between humans and chimpanzees were obtained using previously published syntenic data between human (T2T-CHM13v1.0) and chimpanzee (PanTro6) references ^36^ intersected with SD98 genes using BEDTools intersect. For each paralog, family status was designated as “Human-duplicated gene family” if it was assigned to a gene family containing at least one expanded or duplicated member according to famCN and/or at least one non-syntenic member based on human/chimpanzee synteny. Otherwise, family status was considered “Undetermined”.

#### Paralog-specific copy number genotyping

parCN estimates were obtained using QuicK-mer2 ^47^ for 1KGP 30× high-coverage Illumina individuals ^55^ and four archaic genomes (including Altai Neanderthal [PRJEB1265] ^48^, Vindija Neanderthal [PRJEB21157] ^49^, Mezmaiskaya Neanderthal [PRJEB1757] ^48,49^, and Denisova [PRJEB3092] ^50^), using T2T-CHM13 (v1.0) as reference ^35^. The resulting BED files containing parCN estimates were converted into bed9 format using a custom Python script for visualization in the UCSC Genome Browser. parCN values were genotyped across SD98 regions overlapping protein-encoding and unprocessed pseudogenes by calculating the mean parCN across the region of interest for each sample using a custom Python script.

#### Metrics of selective constraint

Loss-of-function intolerance of SD98 genes was assayed using previously published gnomAD (v2.1.1) probability of loss-of-function intolerance scores (pLI) ^182^ and loss-of-function observed/expected upper fraction (LOEUF) ^53^. We considered genes as intolerant to loss of function if either their pLI scores were greater than 0.9 or their LOEUF scores were less than 0.35.

#### Assessment of variant-calling performance

Despite improved representation of duplicated genes in T2T-CHM13, genomic assessment of these regions remains challenging using short-read Illumina data. We assessed if SNVs were depleted across duplicated regions using variants from 1KGP individuals mapped to T2T-CHM13 (v1.0) ^37^, filtering for biallelic SNPs only, using BCFtools view ^183^ using parameters--exclude-types indels and --max-alleles 2. We compared observed values to empirical distributions, obtained by randomly sampling regions of identical size as SD and SD98 regions using bedtools shuffle with-noOverlapping-maxTries 10000 -f 0.1 parameters. Previously published centromeric satellites coordinates ^184^ were also excluded using the flag - excl. We found that duplicated regions are significantly depleted for SNVs in the high-coverage 1KGP dataset compared to unique regions, which do not include SD or centromeric satellites ^184^ (SD98: 11.79; SD: 25.9, unique: 37.49 SNVs/kbp; *p*-value<0.05 empirical distribution) (Figure S1D). The autosomal 2.4 Gbp in T2T-CHM13 accessible for accurate Illumina SNV calling—determined using read depth, mapping quality, and base quality metrics ^37^—includes only 37.95% and 10.86% of SD and SD98, respectively, while 95.64% of unique space is accessible (Figure S1D). In the SD98 regions, only 56 previously-identified SD98 genes, including 48 protein coding and 8 pseudogenes, are accessible (>90%) to short-reads (Table S1A).

To evaluate our ability to detect variants within duplications using short-read sequencing, we compared SNVs discovered using Illumina short-read versus PacBio HiFi long-read data across eight 1KGP individuals included in both the 1KGP and HPRC ^37,81^ (HG01109, HG01243, HG02055, HG02080, HG02145, HG02723, HG03098, and HG03492). Biallelic SNVs were selected using BCFtools view. Concordance between platforms, measured as precision and sensitivity, was obtained with rtg-tools vcfeval ^185^ for autosomal Non-SDs, SDs, and SD98 regions, using PacBio HiFi variants as a truth-set. Short-read accessible regions were obtained from Aganezov et al. ^37^

While no differences in variant density (SNV sites within 1-kbp non-overlapping windows) existed between technologies in non-duplicated regions, we observed reduced mean variant density from short-read versus long-read data across SD (SRS: 1; LRS: 5) and SD98 (SRS: 0; LRS: 5) (Figure S1E).

Notably, no differences were observed when considering only short-read accessible regions ^37^. Using cHiFi-discovered variants as truth, we next assessed variant calling precision and found that 99.5% of SNVs matched between technologies in non-SD, but decreased to 88.6% and 81.7 % in SD and SD98, respectively (Figure S1F). When considering only short-read accessible regions, SNV precision increased in the three regions assayed to 99.7%, 96.1%, and 94.2% for non-SD, SD, and SD98. Sensitivity— measured as the proportion of HiFi-discovered SNVs also detected using Illumina data—experienced a pronounced decrease of 24.5% in SD and 0.85% in SD98 compared to 87.6% in Non-SD regions. When considering only short-read accessible regions, however, sensitivity is improved to 72.5%, and 57.8% in SD and SD98, respectively. Overall, these results indicate that existing variants identified across duplicated regions from Illumina data are generally accurate, particularly in defined accessible regions, but not comprehensive.

#### Tajima’s D analysis

Additionally, Tajima’s D ^56^ values were calculated using previously published SNPs obtained from high-coverage short-read sequencing data from unrelated 1KGP individuals (n=2,504) ^55^, remapped to T2T-CHM13 (v1.0) ^37^. For each 1KGP continental superpopulation (African, European, East Asian, South Asian, and American), we computed Tajima’s D in 25-kbp windows (≥5 SNPs) using VCFtools ^186^, restricting analyses to short-read–accessible windows (≥50% of bases annotated as accessible in the combined accessibility mask) ^37^. Duplicated and non-duplicated genomic loci differ in several ways, including constraint ^187^ and mutation rates ^154^. To mitigate the effects of these differences, outlier D values were defined as the lower 5th and upper 95th percentiles within accessible SD98 windows (i.e., windows overlapping at least 10% of their bases with SD98 regions and meeting the short-read accessibility criterion). Outlier lower and upper threshold values for each population were defined as follows: AFR, - 2.21 and −0.67; EUR, −2.37 and 0.08; EAS, −2.48 and −0.10; SAS, −2.40 and −0.28; and AMR, −2.40 and - 0.41.

#### Association with neural traits

Due to difficulties mapping short reads to highly identical regions, as well as lack of SD representation on SNP arrays, variants across SD98 genes and regions are depleted in existing genome-wide studies of phenotypes and diseases. To quantify this underrepresentation, we considered the GWAS catalog v1.0 ^188^ (mapped to GRCh38.p12), ClinVar ^189^ (rel. 20200310), and GTEx ^190^ v8 single-tissue eQTL (dbGaP Accession phs000424.v8.p2; mapped to GRCh38, excluding chromosome Y) 12. From the GWAS catalog, we selected only SNPs significantly associated with brain measurements (*p*-value < 0.05) and identified GWAS “mapped genes” that overlapped with our SD98 gene list using gene symbols. We observed significant depletion across the GWAS catalog (SD98: 0.29 variants/100kbp; genome-wide: 1.5 variants/100 kbp), ClinVar (SD98: 20.81 variants/100 kbp; genome-wide: 9.95 variants/100 kbp), and GTEx expression quantitative trait loci (eQTL) databases (SD98: 398.7 variants/100 kbp; genome-wide: 70.14 variants/100 kbp) (Figure S1J).

Additionally, we downloaded published associations between CNVs and neural traits in the UKBB ^58^. Coordinates of CNVs significantly associated with brain measurements (*p*-value < 0.05) were lifted over from hg19 to hg38 and from hg38 to T2T-CHM13 (v1.0) using UCSC liftOver tool ^180^. Liftover chains were obtained from the UCSC Genome Browser and T2T-CHM13 GitHub page (https://github.com/marbl/CHM13, previous assembly releases of T2T-CHM13), respectively. CNVs were intersected with SD98 gene coordinates using BEDTools intersect^179^.

ParCN values from SD98 genes for families with autistic children from the SSC (n = 2,459 families, n = 9,068 individuals) mapped to the T2T-CH13v1.1 reference genome were obtained, following the same steps as described to genotype parCN across 1KGP individuals. Overall, CN differences between autistic probands and unaffected siblings were compared by rounding median CN per individual to the nearest integer, and significance was assessed using the Wilcoxon signed-rank test, correcting for multiple testing with the false discovery rate method. To identify *de novo* deletions or duplications in autistic probands and unaffected siblings, parCN values within ±0.2 of an integer were conservatively selected and rounded to the nearest integer for all family members. Intermediate values, which could potentially confound the analysis, were removed. *De novo* events were classified as cases where both parents exhibited a parCN=2, while the child showed a parCN=3 (duplication) or parCN=1 (deletion).

Previously published genomic hotspots ^191^ were obtained in hg19 coordinates and lifted over to hg38 and from hg38 to T2T-CHM13 (v1.0) using the UCSC liftOver tool and associated chain files (described above). Three regions failed the liftover process due to differences in reference genome sequences. An extra 500 kbp were added upstream and downstream of each reported genomic hotspot to account for breakpoint errors. SD98 genes, including those exhibiting putative *de novo* events in the SSC dataset, were intersected with expanded genomic hotspots coordinates using BEDTools intersect.

#### Gene expression analysis

Previously published brain transcriptomic datasets, including post-mortem tissue and cell lines, were obtained. These datasets included neocortical germinal zones ^62^, neural stem and progenitor cells ^22^, a neuroblastoma cell line SHSY5Y ^63^, and two longitudinal studies of *in vitro* induced neurogenesis from human embryonic stem cells ^64^ (CORTECON), and post-mortem brain (BrainSpan) ^65^—the latter of which was separated into prenatal and postnatal samples. Transcriptomic data from lymphoblastoid cell lines from 69 individuals were also included for comparison ^66^. Raw reads were pseudo-mapped to T2T-CHM13 (v2.0) CAT/Liftoff transcriptome and the CHM13v2.0 assembly as decoy sequence using Salmon v1.8.0 ^192^ with the flags “--validateMappings--gcBias”. The CAT/Liftoff transcriptome was converted to fasta format using gffread ^193^. Transcripts per million (TPM) values and raw counts were summed to the gene level using tximport ^194^. An SD98 gene was considered expressed during development if TPM values were greater than one in at least one of these samples, excluding postnatal BrainSpan data. Conversely, an SD98 gene was considered expressed postnatally if TPM values were greater than one in at least one postnatal stage of BrainSpan.

#### Weighted gene co-expression analysis

WGCNA was performed using the R package ^67^ with two longitudinal brain datasets: prenatal BrainSpan ^65^ and CORTECON datasets ^64^. First, we modeled gene expression across the developmental brain using BrainSpan samples available in NCBI BioProject PRJNA242448. The BrainSpan dataset includes 607 samples from 25 brain regions, spanning early prenatal to adulthood developmental epochs. We filtered for high-quality samples used in a previous gene co-expression analysis ^195^ and focused exclusively on prenatal samples from the frontal cortex, including the dorsolateral prefrontal cortex (DFC; n = 12), ventrolateral prefrontal cortex (VFC; n = 1), medial prefrontal cortex (MFC; n = 12), and the orbital prefrontal cortex (OFC; *n* = 12) (Data S1). Overall, our dataset included 47 samples from four brain regions, spanning three developmental epochs—early prenatal, early-mid prenatal, and late-mid prenatal—covering post-conception weeks (PCW) 8 to 22. We selected genes expressed (≥1 TPM) in at least 80% of the samples for each developmental epoch and brain region, resulting in 17,388 genes which were used as input for WGCNA analysis. We used Principal Component Analysis (prcomp function in R) to cluster samples based on their expression profiles, and no outliers were identified (Data S1). However, the first two principal components distinctly separated the early prenatal samples at 8 weeks post-conception (8 PCW) from the rest, indicating that post-conception age exerts a stronger influence on gene expression than the broader developmental epoch.

To overcome coverage and batch effects, we performed variance stabilizing transformation on the raw counts using function varianceStabilizingTransformation() from the R package DESeq2 with a blind design to preserve variability within each developmental epoch. We use the function pickSoftThreshold() from the R package WGCNA to estimate the power parameter that generates a scale-free topology network, choosing the minimum value where the scale-free topology fit index is around 0.8, which in the BrainSpan dataset was 24. Normalized counts were used as input to construct a signed network using function blockwiseModules() with parameters networkType=“signed”, deepSplit=4, detectCutHeight=0.995, minModuleSize=30, mergeCutHeight=0.25, and softPower=24, and default parameters otherwise. This yielded 23 modules represented by their respective eigengenes (Data S1) named “B-” followed by a random color, ranging from 47 to 3,126 genes each, with a median module size of 428. The largest modules were B-turquoise (n=3,126), B-blue (n=2,727), B-brown (n=2,102), and B-yellow (n=1,802). Despite the larger numbers of genes assigned with these clusters, the median module memberships remained high (B-turquoise: 0.72, B-blue: 0.74, B-brown: 0.73, B-yellow: 0.72). The B-turquoise and B-yellow were negatively correlated with early prenatal stages, while module B-blue and B-brown were correlated with prenatal stages. Seventeen modules included paralogs from human-specific gene families. In the BrainSpan dataset, 2,320 genes remained unclustered and were assigned to module B-Grey, reflecting the expression noise inherent in complex tissues composed of multiple cell types.

We then modeled gene expression using the CORTECON dataset, which tracks *ex-vivo*-induced neurogenesis from human embryonic stem cells. This dataset includes 23 transcriptome samples (our analysis excluded the test set), spanning the stages of pluripotency, neural differentiation, cortical specification, deep-layer formation, and upper-layer formation (Data S1. We clustered samples based on their raw expression counts using hierarchical clustering and Principal Component Analysis (prcomp function in R), which flagged two samples as outliers (SRR1238515 and SRR1238516) and therefore they were removed from downstream analyses. To reduce noise, we removed gene features with consistently low counts, meaning less than 10 counts across 90% of the CORTECON samples, resulting in 15,697 gene features. We transformed raw counts using the varianceStabilizingTransformation() function from the DESeq2 package with a non-blind design that aimed to remove differences from developmental stages. To determine the optimal softPower parameter for the CORTECON dataset, we used the pickSoftThreshold() function from the WGCNA package, which identified a value of 24.

Co-expressed modules were obtained with the function blockwiseModules() from the WGCNA package, using networkType=”signed”, deepSplit=4, detectCutHeight = 0.995, minModuleSize=30, mergeCutHeight = 0.15, and softPower=24, and default parameters otherwise. This approach yielded 37 co-expressed modules (Data S1), represented by their respective eigengenes and named as “C-” followed by a random color. The number of genes assigned to a module ranged from 33 to 3,988, with a median value of 424.1 genes per module. The most genes were assigned to modules C-turquoise (n=3,988), C-blue (n=2433), C-yellow (n=1,149), C-green (n=954), and C-red (n=709). The median module memberships for these larger modules were higher than the BrainSpan dataset, likely reflecting reduced variability in the *in vitro* neuronal samples versus the more heterogeneous post-mortem brain tissue (C-turquoise: 0.81, C-blue: 0.84, C-yellow: 0.85, C-green: 0.86, and C-red: 0.83). These modules showed the strongest associations with developmental stages, where C-turquoise and C-green were strongly associated with pluripotency (Pearson r = 0.87 and 0.97, respectively), C-yellow and C-blue were strongly anti-correlated with pluripotency (Pearson r = −0.98 and −0.9, respectively), and C-red was correlated with neural differentiation and anti-correlated with upper layer formation (Pearson r = 0.67 and −0.56). Twenty-one co-expression modules included paralogs from human-specific gene families. Importantly, 871 genes were assigned to the C-grey module, which corresponds to unclustered genes (Data S1). To verify CORTECON module assignments, we also assessed human-duplicated genes with known functions. *ARHGAP11B*, which induces cortical neural progenitor amplification by altering glutaminolysis in the mitochondria ^24^, is a member of the C-turquoise module. Genes in this module are expressed highest during pluripotency and are associated with cell proliferation, including DNA replication and chromosome segregation, as well as mitochondrial gene expression. Additionally, the hominoid-specific gene *TBC1D3*, known to promote basal progenitor amplification in the outer radial glia resulting in cortical folding in mice ^25^ is a member of the C-purple module, which is associated with regulation of neural differentiation.

Module concordance was calculated for each gene family as the proportion of its members assigned to the same module, defined as the maximum number of co-assigned members divided by the total number of members in the family. A concordance score of 1 indicates that all members were assigned to the same module, while a score of 0 indicates that no members shared a module assignment. Visualization of the yellow network was constructed by selecting genes with module membership greater than 0.5, generating an adjacency matrix with remaining genes, and then reconstructing a signed network with soft threshold = 18. Edges with Pearson correlation <0.1 were removed. The network visualization was built with the igraph R package (https://r.igraph.org/), using layout_with_fr for vertex placement. Vertex size was proportional to the degree and edges width was proportional to the Pearson’s correlation coefficient. Some vertices were manually adjusted to improve aesthetics of the plot. GO terms enrichment analysis was performed using the R package clusterProfiler ego function, using an adjusted *p*-value threshold of 0.05 ^196^. Enrichment of gene categories were performed using the hypergeometric test in R for autism genes ^68^, expanded genomic hotspots ^191^, and cell markers ^195^, as well as for SD98 genes and human duplicated genes.

#### Mouse and zebrafish orthologs

Mouse-human orthologs were obtained from the Mouse Genome Informatics (MGI) complete list of human and mouse homologs and ENSEMBL BioMart, intersected with SD98 genes using gene symbols, and manual curation. Zebrafish-human orthologs were obtained from combined ENSEMBL BioMart annotations, MGI complete list of vertebrate homology classes, and manual curation. MGI files were downloaded from their website (https://www.informatics.jax.org/homology.shtml) and BioMart analyses were performed using the R package biomaRt. Comparison of developmental brain expression of SD98 orthologs in model organisms was performed using previously published expression data for mouse (PRJNA637987) ^75^ and zebrafish (GSE158142) ^76^, calculating Z-score normalized TPM values. Matching of developmental stages across human, mouse, and zebrafish was done as previously described ^77^. In brief, genes with one-to-one orthologs with human genes were identified (mouse n= 19,949; zebrafish n= 16,910) and the principle component analysis rotations of the human BrainSpan data used to predict PC coordinates for the mouse and zebrafish data in human principle component space.

#### Capture HiFi sequencing

We performed cHiFi sequencing of 172 individuals from the 1KGP, two trios from Genome in a Bottle ^174^, and 22 HGDP individuals with available linked-read data via the 10X Genomics platform ^173^, totaling 200 samples and 18 family trios (Table S4B). DNA samples for 1KGP and Genome in a Bottle were obtained from the Coriell Institute (Camden, NJ, USA) and HGDP samples were obtained from the CEPH Biobank at the Fondation Jean Dausset-CEPH (Paris, France). PacBio cHiFi sequencing was performed using the RenSeq protocol ^197^. Briefly, genomic DNA (∼4 μg) was sheared to approximately 3 kbp with the Covaris E220 sonicator using Covaris blue miniTUBEs, followed by purification and size selection with AMPure XP beads. End repair and adapter ligation were performed using the NEBNext Ultra DNA Library Prep Kit. Barcodes to distinguish each sample were added via PCR using Kapa HiFi Polymerase (Roche, CA, USA). After the first PCR (fewer than 9 cycles), the libraries were purified and size-selected. For target enrichment, 80-mer RNA baits were designed and tiled at 2× coverage across targeted SD regions and unique exonic regions (Table S4D). pHSD regions of interest were targeted and enriched for using a custom myBaits kit (Arbor Biosciences, MI, USA) following manufacturer’s recommended protocol. Eight pooled barcoded libraries were hybridized overnight to the baits, and the captured DNA was bound to Dynabeads MyOne Streptavidin C1 beads. A second PCR was performed post-hybridization to generate sufficient material for sequencing. A PCR cycle test was conducted prior to the second amplification to limit PCR duplication bias.

The final libraries were size-selected using the Blue Pippin system to enrich for fragments >2 kbp and sequenced on the PacBio Sequel II platform (Maryland Genomics, University of Maryland). Briefly, Sequel II libraries were constructed using SMRTbell Express Template Prep Kit 2.0 (Pacific Biosciences, Menlo Park, CA) according to manufacturer’s instructions. In brief, DNA samples were treated with DNA-damage repair enzymes followed by end-repair enzymes before being ligated to overhang sequencing adaptors. Libraries were then purified with SPRI beads (Beckman Coulter, Indianapolis, IN) and quantified on the Femto Pulse instrument (Agilent Technologies, Santa Clara, CA). Prior to sequencing, libraries were bound to Sequel II polymerase, then sequenced with Sequel II Sequencing kit and SMRT cell 8M on the Sequel II instrument (Pacific Biosciences, Menlo Park, CA).

The capture sequencing protocol included tiled baits across all duplicated regions of interest and only exons in non-duplicated space (Figure S4B). As a result, unique exons exhibited significantly lower coverage compared to duplicated exons (Mann-Whitney U test, *p*-value=2.2×10^-16^). Importantly, we did not observe significant differences in coverage between ancestral and derived paralogs, despite the baits being designed based on the ancestral sequence (Mann-Whitney U test, *p*-value>0.05). cHiFi coverage across regions of interest was calculated using samtools depth ^183^ with--min-MQ 10. Globally, considering a cutoff MAPQ score greater than 10, we achieved ∼3 kbp reads with an average coverage of 27× within regions of interest (Table S4E, Figure S4C). We also assessed for the occurrence of PCR duplicates given that they pose three problems: 1) the true output of diverse representation of reads that are sequenced is reduced, 2) lead to false positive variant calls skewing allele frequencies, and 3) may introduce erroneous mutations that do not reflect true population variants. We found 66% of sequenced reads to be unique genome-wide, and within the intended capture space, 34% of the total unique reads mapped to the regions of interest.

#### Long-read genetic variation

Fully phased haplotypes from 47 individuals from the HPRC Year 1 freeze (https://github.com/human-pangenomics/HPP_Year1_Data_Freeze_v1.0) and 15 from the HGSVC ^84^ were downloaded. Each haplotype was mapped to T2T-CHM13v1.0 reference genome using minimap2 with parameters-a--eqx - -cs -x asm5 --secondary=no -s 25000 -K 8G, and unmapped contigs and non-primary alignments were discarded. For each region of interest, the longest alignment spanning the locus was selected and additional alignments were removed. This process ensured that one single contiguous contig was used for variant detection. Variants were called with htsbox ^198^ pileup with parameters -q 0 -evcf and converted into diploid calls using dipcall-aux.js ^199^ vcfpair. For each region of interest, individual sample calls were merged into a multi-sample VCF file using BCFtools merge, only including individuals whose two haplotypes fully spanned the region of interest. Redundant samples between the HPRC and HGSVC (HG00733, HG02818, HG03486, NA19240, NA24385) were removed, prioritizing HPRC assemblies.

cHiFi reads were processed using the standard PacBio SMRT sequencing software tools available in the Conda repository pbbioconda. Circular consensus was obtained from subreads using CCS command with the following parameters --minPasses 3 and --minPredictedAccuracy 0.9. PacBio adapters and sample barcodes were removed using lima software and duplicates were removed with pbmarkdup. Resulting cHiFi reads were aligned to T2T-CHM13v1.0 reference using pbmm2 align, a wrapper of minimap2, with the CCS preset and default parameters. For each sample, read groups were added with Picard AddOrReplaceReadGroups and variants were called on each sample using GATK HaplotypeCaller ^200^, using ploidy = 2 and minimum mapping quality thresholds for genotyping of 0, 2, 5, 10 and 20, resulting in gVCF files per sample for joint genotyping. Joint genotyping was performed with GATK CombineGVCFs and GenotypeGVCFs tools using the pedigree file for accurate calculation of inbreeding coefficients. Genotyping was performed using minimum genotyping confidence thresholds of 0, 10, 20 and 30, and variants were subsequently filtered using hard-filtering thresholds for both genotyping quality (0, 20, 50, 70) and depth (0, 4, 8, 12, 16).

We optimized variant genotyping and hard-filtering parameters by benchmarking minimum thresholds using both population and trio-based analysis, focusing exclusively on biallelic SNPs selected with bcftools view--max-alleles 2 and bcftools view--exclude-types indels. Specifically, we assessed deviations from Hardy-Weinberg equilibrium calculating inbreeding coefficients from the founder population (excluding offspring), with an inbreeding coefficient below −0.3 considered indicative of excess heterozygosity. Additionally, we evaluated Mendelian concordance within trios, calculated for each threshold combination using rtg mendelian ^185^, excluding trios where any of the members had a missing genotype with bcftools view -i ‘F_MISSING=0’. Total number of variant sites were obtained with BCFtools stats.

As MAPQ and genotyping confidence thresholds became more stringent, the total number of variant sites decreased, while biological metrics improved, including increased Mendelian concordance and reduced excess heterozygosity (Figure S4D,E). A minimum MAPQ threshold of 20 reduced sites with excess heterozygosity and improved Mendelian concordance across all genotyping confidence levels, while only marginally reducing the number of detected variants. Therefore, we conservatively proceeded with a minimum MAPQ of 20 and a minimum genotyping confidence threshold of 30. We next optimized hard-filtering parameters, observing that Mendelian concordance increased significantly with higher read-depth and genotype-quality thresholds. We achieved near 100% concordance using either genotype quality of 50 (at any read depth) or the combination of read depth 8 with genotype quality 20. Since the latter combination provided similar performance while retaining more variants, we selected these as our hard-filtering parameters, resulting in the identification of 28,476 biallelic SNVs across 200 individuals. For downstream population genetics analyses, we retained the 144 unrelated individuals that passed our quality thresholds and merged their cHiFi variants with variants from 56 non-redundant HPRC/HGSVC individuals using BCFtools merge, creating a unified cohort of 200 genomes. Functional consequences in the combined dataset were annotated with the Ensembl Variant Effect Predictor (VEP).

Haplotype networks for *CD8B* were constructed using HPRC/HGSVC continuous haplotypes extracted with BEDtools getfasta and aligned with Muscle using Mega Software ^201^. Networks were generated using a minimum spanning tree with the software PopArt ^202^.

#### Tests for signatures of natural selection

Ka/Ks ratios (also known as dN/dS) were calculated for pHSD paralogs, performing pairwise comparison between human and chimpanzee sequences, based on T2T-CHM13v1.0 and panTro6 reference genomes, respectively. Alignments between human and chimpanzee canonical transcripts sequences were manually curated and used as input for seqinr package for Ka/Ks estimation. pN/pS ratios were calculated using as input variant sites estimated by seqinr package as well as polymorphic variation from the combined cHiFi and HPRC/HGSCV dataset, considering only biallelic SNPs from unrelated samples (n=144).

Synonymous and nonsynonymous mutations were defined based on previously calculated VEP consequences. Ka/Ks and pN/pS values were jointly analyzed using the Direction of Selection (DoS) statistic, a derivation of McDonald–Kreitman’s neutrality index, defined as DoS = Dn/(Dn + Ds) - Pn/(Pn + Ps) ^88^. Significant differences in Ka/Ks or DoS between ancestral and derived paralogs were assessed using Wilcoxon signed-rank test, pairing each derived paralog to its ancestral counterpart. dN/dS was determined, in parallel, across gene families using codeml as part of the Phylogenetic Analysis by Maximum Likelihood (PAML ^89^) from generated multiple-species alignments for each gene family (MAFFT ^203^), using T2T-CHM13 for human paralog sequences and orthologous sequences from respective genomes for chimpanzee (panTro6), gorilla (gorGor6), orangutan (ponAbe3), rhesus (rheMac10), mouse (mm39), and rat (rn7). Ancestral and derived states for pHSD genes were assigned based on previously published predicted states ^14^. Conservatively, the evolutionary status of four gene families was considered as “unknown” and excluded from calculations of statistical differences (*FRMPD2*/*FRMPD2B*, *PTPN20*/*PTPN20CP*, *GPRIN2*/*GPRIN2B*, and *NPY4R*/*NPY4R2*). Paralogs with infinite values were also excluded from the analysis.

Nucleotide diversity (π) and Tajima’s D statistics were calculated across selected pHSD loci using biallelic SNPs derived from continuous haplotypes from HPRC and HGSVC assemblies, utilizing the PopGenome ^204^ R package and its functions F_ST.stats and neutrality.stats, respectively. For the gene bodies of *GPR89*, *ROCK1*, *FAM72*, and *CD8B*, π and Tajima’s D values were calculated using 15-kbp windows with 1-kbp steps. For *GPR89* paralogs, π was calculated across extended surrounding duplicated regions using 20-kbp windows and 1-kbp steps. For *CD8B* paralogs, Tajima’s D was calculated in surrounding regions using 6-kbp windows and 500-bp steps.

#### Generation of zebrafish lines

Creation of CRISPR lines to knockout genes of interest was done as previously described ^105,109,205^. Briefly, crRNAs were annealed with tracrRNA (Alt-R system, Integrated DNA Technologies, Newark, NJ) in a 100 µM final concentration to make the sgRNA duplex, which was then coupled with SpCas9 (20 µM, New England BioLabs, Ipswich, MA) to prepare injection mixes. All oligonucleotide sequences can be found in Table S5A. Microinjection of one-cell stage zebrafish embryos was performed using an air injector (Pneumatic MPP1-2 Pressure Injector) to release ∼1 nl of injection mix into each embryo.

Injection mixes to knockout-specific genes included ribonucleoproteins with four different sgRNAs targeting early exons in equimolar concentrations. In parallel, stable CRISPR knockout lines were made using a single sgRNA (Table S5A). Knockout alleles in stable lines corresponded to a 5-bp deletion in *frmpd2* (named *frmpd2^tupΔ^*^5^) and an 8-bp deletion in *gpr89* (named *gpr89^tupΔ^*^8^, allele sequences can be found in Table S6E). For *arhgap11* knockdown, morpholinos blocking translation (GeneTools, Philomath, OR) were reconstituted to 2 mM and ∼1 nl of a 2 ng/nl mix was microinjected into one-cell-stage embryos. Assessments of potential off-target sites for all sgRNAs used in this study were performed with the CIRCLE-seq protocol ^206,207^ and top potential off-target sites were evaluated via Sanger sequencing as previously described ^105^. No editing was observed in potential off-target sites for any sgRNA used in this study, suggesting that phenotypes observed are due to the targeted knockout.

“Humanized” zebrafish larvae were generated by temporal expression of transcribed mRNAs. Expression vectors containing human transcripts were used to generate mRNA, including pEF-DEST51 (*SRGAP2C* and *ARHGAP11B*), pGCS1 (*GPR89B*, *PDZK1P1*, and *PTPN20CP*), pCR-TOPO (*NPY4R* and *FAM72B*), and pCMV-SPORT6 (*FRMPD2B*). The cDNA inserts of two genes were synthesized (Twist Biosciences, San Francisco, CA) based on transcript evidence from IsoSeq data from the ENCODE ^208^ project (*PDZK1P1*: ENCFF158KCA, ENCFF939EUU; *PTPN20CP*: ENCFF305AFY). All plasmids were sequenced through either Azenta or Plasmidsaurus. Following plasmid linearization using restriction enzyme digest and DNA purification, 5’-capped *in vitro* mRNA was generated using the MEGAshortscript transcription kit (Thermo Fisher, Waltham, MA) following the manufacturer’s protocol with a 3.5 h 56°C incubation with T7 or SP6 RNA polymerase, depending on the plasmid. The resulting transcripts were purified with the MEGAclear transcription clean-up kit (Thermo Fisher, Waltham, MA), measured quantity with the Qubit, and visualized on a 2% agarose gel to ensure intact transcript. All mRNA injection mixes included mRNA at a 100 ng/μl concentration and ∼1 nl of the mix microinjected into one-cell stage embryos, as described above. Presence of the human mRNA transcripts was observed by using 500 ng of extracted RNA from 3 dpf-injected larvae used in the sciRNA-seq experiment with the SuperScript IV Reverse Transcriptase kit (Thermo Fisher, Waltham, MA) followed by PCR amplification with DreamTaq PCR Master Mix (Thermo Fisher, Waltham, MA) and primers listed in Table S5A.

#### Morphometric assessments

High-throughput imaging of the zebrafish larvae was performed using the VAST BioImager system (Union Biometrica, Holliston, MA) as previously described ^108,109^. Mutant and control larvae at 3 or 5 dpf were placed into 96-well plates where they were then acquired by a robotic arm, placing the larvae in a rotating 600 µm capillary coupled with a camera, allowing for the automatic acquisition of images from four sides. Images were then processed and analyzed using the TableCreator tool in FishInspector v1.7 ^107^ to measure the head area and body length of 3,146 larvae—discarding images with general issues (e.g., dead or truncated larvae). To validate changes in head area, a neuronal reporter transgenic zebrafish line Tg[HuC-GFP] ^121^ was used to create CRISPR-knockouts or humanized larvae that were then kept in an incubator at 28°C until imaged at 3 dpf using tricaine as anesthesia (0.0125%) and low-melting agarose. Imaging was performed in the Dragonfly spinning disk confocal microscope system with an iXon camera (Andor Technology, Belfast, United Kingdom). Z-stacks of 10 µm slices for each larva were collected and processed using Fiji ^209^ to generate hyperstacks with maximum intensity projections. Forebrain areas were measured in a blinded manner by a different trained investigator by manually delimiting the forebrain region. Any image with tilted larvae or unclear definition of the different brain regions was not included.

#### Supervised classification of images

As an alternative to performing statistical tests to identify changes in predefined morphological measurements between mutants and controls, we employed a convolutional neural network (CNN) to identify differences between mutants and controls without the need to measure predefined features ^110^. Due to the use of multiple 96-well plates for each mutant, we observed significant batch effects in the resulting images, where larvae images from the same plate were significantly more similar to each other than to genotypically matched larvae from different plates. Therefore, before training our CNN-based classifier, we trained a latent diffusion model (LDM) to minimize the plate batch effect before input into the CNN. The goal of the LDM is to use the larvae with control genotypes present on each plate to learn the plate-specific batch effects, or style. We then select a single plate as a reference and use the LDM to transform all images to the reference plate style, therefore making them comparable. The LDM removes batch effects by first transforming the original image *x* into a latent representation *z_0_* through a variational autoencoder function *ε* that reduces the dimensionality of *x* but does not remove any batch effect:

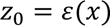

We then pass the encoded image *z_0_* through an LDM forward process, in which we repeatedly sample new latent variables *z_1_*,…,*z_T_* by effectively repeatedly adding Gaussian noise to the original image *z_0_*:

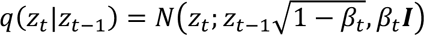

Where *β_t_* is the magnitude of noise decided by a noise scheduler at time step *t.* This ultimately transforms the original (VAE-transformed) image *z_0_* into the embedding *z_T_*. This embedding *z_T_* represents the larvae as an embedded image, free of association with any batch effect.

In the third step, we apply a reverse process of the LDM by successively transforming the image *z_T_* into a new *z_0_,* but ‘add back in’ the effect of a reference plate batch by introducing a condition variable *c* which comprises the desired batch and mutant ID. This conditional reverse process can be expressed as:

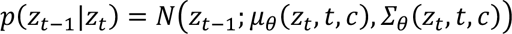

Where *μ_θ_(z_t_,t,c)* and *Σ_θ_(z_t_,t,c)* represents the predicted mean and covariance functions. Finally, we pass our model through the decoder of a variational autoencoder to reconstruct the original image *x* into a new image, *x’*, that represents the original image *x* but in the new reference plate style, suitable for input into the CNN classifier.

Our LDM is trained to minimize negative log likelihood using 350 diffusion steps with a linear noise scheduler ^210,211^. After training the model, we applied the model to transform all images to one reference plate, which is selected as the one with the highest number of controls. This transformation process minimizes the batch effect by generating images that appear as if they were collected from the same plate.

Having minimized batch effects on the larvae images, we then trained a CNN image classifier to determine the extent to which each mutant genotype differs from matched controls on the basis of the raw morphometric images alone. Higher classification accuracy, as measured by F1 score, indicates a larger effect size of mutant genotype on morphology. Our CNN framework involved fine-tuning a pretrained Alexnet classifier on the transformed larvae images ^212^. More specifically, we trained 17 different Alexnet classifiers, one per mutant genotype, to perform binary classification to distinguish one specific mutant genotype from controls. The models were trained and evaluated in a five-fold cross validation framework, with F1 scores averaged over all folds. To generate feature attribution heat maps highlighting the morphological regions used to distinguish each mutant genotype from controls (Figure 5B), we used the GradCAM (Gradient-weighted Class Activation Mapping) approach ^165^. We selected a *GPR89B* and *gpr89KO* sample representative of the pattern exhibited across mutants from this family. All code related to this analysis is available ^213^.

#### Single-cell RNA-seq

We performed cellular assessments using the single-cell combinatorial indexing RNA sequencing (sciRNA-seq) protocol ^113^. Zebrafish larvae from CRISPR knockout or mRNA-injected lines were generated as described above and kept in an incubator at 28°C until 3 dpf when they were euthanized in cold tricaine (0.025%) and their heads immediately dissected, pooling 15 heads together per sample.

Dissociation of the dissected heads was performed following two washes in 1 ml of cold 1x PBS on ice with a 15 min incubation in dissociation mix (480 µl of 0.25% trypsin-EDTA and 20 µl of collagenase P at 100 mg/ml), gently pipetting each sample every 5 min with a cut-open P1000 tip for complete dissociation. Once all tissue was visibly dissociated, 800 µl of DMEM with 10% FBS was added to each sample and centrifuged for 5 min at 700g at 4°C, resuspended in cold 1x PBS and centrifuged again at 700g for 5 min at 4°C. Cells were then resuspended in 800 µl of DMEM with 10% FBS and filtered through a 40 µm cell strainer (Flowmi, Sigma Aldrich, St. Louis, MO) using low-bind DNA tubes (Eppendorf, Hamburg, Germany). Cells were counted using a Countess II (Thermo Fisher, Waltham, MA) and all samples with viability >65% used further. Immediately after viability confirmation, cells were fixed as previously described ^214^ with a 10 min incubation in 1.33% formaldehyde in 1x PBS on ice followed by permeabilization with 5% Triton-X for 3 min on ice, and neutralization with 10% Tris-HCl (1M, pH 8). Cells were then filtered through a 40 µm cell strainer again, 15 µl of DMSO added to each sample in 5-µl increments, and then slowly frozen in a Mr. Frosty (Thermo Fisher, Waltham, MA) freezing container filled with isopropanol at −80°C overnight.

Library preparation was performed following the sciRNA-seq protocol as described ^113^, including three rounds of combinatorial indexing of the cells (all primer sequences correspond to Plate 1 of the original protocol and can be found in www.github.com/JunyueC/sci-RNA-seq3_pipeline). The first round involved reverse transcription with barcoded oligo-dT primers to introduce the initial index. Cells were then pooled and redistributed into new wells for the second round, where a second index was added via ligation. The third round included second-strand synthesis, tagmentation with Tn5 transposase, and PCR amplification to incorporate the final index. Libraries were evaluated for quality control in a BioAnalyzer and Qubit to check integrity and concentration, and then sequenced in three NovaSeq 6000 lanes (Novogene, Sacramento, CA). Raw fastq files were processed following the available sci-RNA-seq3 pipeline ^215^ (www.github.com/JunyueC/sci-RNA-seq3_pipeline). This pipeline includes attachment of the unique molecular identifier (UMI) sequence to each read2 based on the identified RT and ligation barcodes from read1 (edit distance ≤1), and trimming with TrimGalore v0.4.1 (https://zenodo.org/records/7598955), using cutadapt ^216^ and fastqc ^217^. Reads were then mapped to the improved zebrafish transcriptome ^218^ with STAR ^219^ using the--outSAMstrandField intronMotif option. Duplicates (reads with the same UMI) were removed with the available custom-made python scripts found in the Cao lab GitHub repository. Lastly, filtered SAM files were split by their UMI sequences (corresponding to individual cells) and gene-cell count matrices constructed by mapping reads to the zebrafish v4 GTF file ^218^.

Gene-cell count matrices were loaded into R to generate Seurat v4 ^220^ objects and cells with transcript counts below 150 or above two standard deviations over the mean, mitochondrial or ribosomal gene counts >5%, or potential doublets (with a ∼4% doublet expectation based on previous reports ^215,221^ and estimated using DoubletFinder ^222^) were removed (Figure S5E). Cells from different libraries were normalized using SCTransform ^223^ with the glmGamPoi method and regressing by the percentage of mitochondrial and ribosomal counts. Then, normalized counts across sequencing libraries were integrated with Harmony ^224^ with a PCA reduction using batch as a grouping variable. Hierarchical clustering was performed by calculating the euclidean distances across all cells using the Harmony cell embeddings and clustering with the hclust function using the ward.D2 method. The hierarchical tree was cut at a K of 50, gene markers for each cluster estimated using the FindAllMarkers function (logfc.threshold=0.10, test.use=”MAST”, min.pct=0.15, min.diff.pct=0.10), and classification into cell types using available zebrafish brain scRNA-seq atlases ^76,119^ and the Zebrafish Information Network (ZFIN ^98^) website.

Focusing on neuronal, glial, and eye-related clusters left a total of 95,555 cells for further analysis (Table S5F). General correlations across samples (knockout vs. “humanized” models for each gene of interest) were done with a balanced number of cells for each pair and pseudo-bulking gene counts by sample and cluster, so counts across cells were summed together for each sample, allowing for biological replicates to be maintained. Then, pseudo-counts were processed with DESeq2 ^225^ with the Wald test option to obtain fold-change values for each gene compared to their respective control (SpCas9-scrambled gRNA injected for crispants, GFP-mRNA-injected for “humanized”, and control-morpholino-injected for *arhgap11*-knockdown). Then, cell-type-specific differential gene expression tests were performed similarly but with previous subsetting of the matrix for each cell type. For *FRMPD2* and *GPR89* models, forebrain cells were further re-clustered to obtain more detailed cell types; gene expression across samples correlated as described above using a pseudo-bulk approach with the telencephalic cells. Progenitor and differentiated cell classification was performed using known neural progenitor (*sox19a*, *sox2*, *rpl5a*, *npm1a*, *s100b*, *dla*) or mature neuron (*elavl3*, *elavl4*, *tubb5*) markers and the PercentageFeatureSet function to estimate the weight of these genes per cell.

#### Seizure susceptibility

To assess changes to chemically induced seizure susceptibility, we employed an optimized published protocol ^128^. Briefly, larvae were collected and kept in an incubator at 28°C until 4 dpf, when they were distributed in a 96-well plate and placed in a Zebrabox system chamber (ViewPoint, Montreal, Canada) that has a camera with an acquisition speed of 30 frames per second. Treatments included 0 or 2.5 mM of pentylenetetrazol (PTZ, #P6500, Sigma-Aldrich, St. Louis, MO) in a total volume of 200 µl per well.

Once placed in the Zebrabox chamber, larvae were left for 10 min unbothered before starting a 15 min recording (acquisition in 1 s bins) to then extract the frequency of high-speed events (>28 mm/s) using a published MATLAB script ^128^ to compare against batch-sibling controls.

### Quantification and statistical analysis

#### Gene ontology overrepresentation

Overrepresentation of gene ontology terms across human duplicated genes and copy-number constrained genes was assessed with clusterProfiler ^196^, using all human-genes as background and considering terms with Benjamini-Hochberg adjusted *p*-value ≤ 0.05 as significant. Enrichment of DEGs in GO terms for the zebrafish pseudo-bulk analysis was also estimated with clusterProfiler ^196^ using only the expressed genes as the background list for the tests and *p*-values below 0.05 after a Benjamini-Hochberg adjustment were considered as significant.

#### Genetic association

To assess depletion levels of associated variants from published databases, observed values were compared to empirical null distributions, built from 10,000 non-overlapping, size-matched random regions generated using bedtools shuffle-noOverlapping-maxTries 10000 -f 0.1. One-tailed empirical *p*-values were calculated as: *p* = (M + 1) / (N + 1), where M is the number of iterations yielding a number of features less than (depletion) the observed value and N is the number of iterations. Empirical *p*-values were calculated using 10,000 permutations. Significant differences between CN across probands and unaffected siblings were calculated using a paired design for each sibling pair, using a Wilcoxon signed-rank test corrected for multiple testing using false discovery rate, considering *q*-values ≤ 0.05 as significant.

#### Brain expression

Significant differences in expression across developmental stages of the CORTECON datasets stratified by copy-number status (polymorphic, nearly fixed and fixed) were obtained using a Mann-Whitney U test. Significance thresholds were defined as: ns: non-significant; * ≤ 0.05, ** ≤ 0.01 and *** ≤ 0.001. Enrichment of gene categories (human duplicated, SD98, autism, human hotspots genes) across BrainSpan and CORTECON modules was assessed with a Hypergeometric test (phyper function in R) using *p*-value ≤ 0.05 as significance cut-off. Gene ontology overrepresentaton analysis across BrainSpan and CORTECON modules was performed with clusterProfiler ^196^, employing all human genes as the background and considering terms with a Benjamini-Hochberg *p*-value ≤ 0.05 as significant.

#### Signatures of natural selection

Outlier genome-wide Tajima’s D values were obtained as the 5th and 95th percentiles across each 1KGP continental population. Differences in heterozygous sites distribution between paralogs across pHSD families were calculated using a Mann-Whitney U test, considering *p*-value ≤ 0.05 as significant.

Differences in allele frequency distributions between 35-kbp windows overlapping *CD8B* and *CD8B2* loci were obtained using Kolmogorov-Smirnov test. Global differences in Ka/Ks and Direction of Selection values between ancestral and derived pHSD genes were obtained using a paired-design between ancestral and derived paralogs, using a Wilcoxon signed-rank test. For gene families with more than two paralogs, all derived paralogs were compared to the ancestral one.

#### Single-cell transcriptomics

To define cellular identities of each obtained cluster, we extracted DEGs between clusters using the MAST test with a Bonferroni correction. For the cell-type differential expression between genotypes, we balanced the number of cells between groups (maintaining the number of cells from the smaller group) and gathered gene counts to obtain pseudo-bulk values that we then compared using the Wald test with a Benjamini-Hochberg adjustment. Fold-change correlations between DEGs across groups was assessed using the spearman method. Evaluation of neuronal classification in progenitor-like or mature was done using a generalized linear model after each neuronal cell was assessed for the presence of counts for the defined progenitor or mature neuronal markers described above. In all analyses, adjusted *p*-values below 0.05 were classified as significant.

#### Morphometrics

For each morphometric measurement, values larger than the 75% quantile plus two times the interquartile range or smaller than the 25% quantile minus two times the interquartile range were considered as technical outliers and removed from the distribution. Normality of each morphometric value was assessed using a qq-plot and a Shapiro-Wilk test. Due to non-normality of residuals, an ANCOVA with a rank-transformation for each measurement was used with the date of the experiment and the plate as covariates. To control for potential confounding technical biases, for each mutant evaluated, we compared all larvae to the controls obtained in the same experimental batch. *p*-values from paired tests between mutants and controls were adjusted using the Benjamini-Hochberg method and significance was determined as adjusted *p*-values below 0.05. All numbers of larvae included in each comparison can be found in Table S5E, as well as each raw and adjusted *p*-values, the mean values for each group, standard deviation, and delta. For the stable zebrafish lines, direct comparisons between mutants and controls were performed using the Wilcoxon test due to non-normality of residuals and no additional covariables were included given that all values were obtained in the same experiment. No multiple testing adjustment was added to these Wilcoxon rank-test *p*-values. The number of larvae included in each group for these tests can be found in Figure 6 at their corresponding boxplots, which include the median and quantile values. The values above each box is the *p*-value from the Wilcoxon tests against controls.

#### Seizure susceptibility

Frequencies of HSE between control and mutant groups was performed using a Kruskal-Wallis test with a Dunn’s post-hoc test given that values were non-normally distributed following an assessment via qq-plot and Shapiro-Wilk tests. The number of larvae included in the reported groups in Figure 6F for the control treatment (0 mM PTZ) were 666 controls, 54 *frmpd2* knockouts and 65 *FRMPD2B*-injected, while the 2.5 mM PTZ treatment included 693 controls, 84 *frmpd2* knockout and 71 *FRMPD2B*-injected larvae. Tests with a *p*-value below 0.05 are highlighted in the heatmap with an asterisk.

## Additional resources

- parCN estimates for 1KGP individuals available at https://dcsoto.shinyapps.io/shinycn.
- Gene expression from human brain datasets available at https://dcsoto.shinyapps.io/shinybrain/.

## Supplemental Information

**Figure S1. Detailed genetic analysis of human duplicated genes related to Figure 1 and STAR Methods. (A**) Pipeline to group SD98 genes into gene families. (**B**) Distribution of number of gene members within duplicate gene families. (**C**) gnomAD pLI versus LOEUF scores for all SD98 genes with available scores. (**D**) From top to bottom: 1KGP short-read SNVs ^37^ in SD (blue, left) and SD98 (orange, right) using the T2T-CHM13 regions. Observed values are shown as vertical bars, while empirical distributions of the number of variants observed in randomly sampled regions are represented as density plots. Total region size (in Gbp) and accessible sites size (darker colors), for NonSD (gray), SD (blue), and SD98 (orange). **(E)** Distribution of biallelic SNVs across non-overlapping 1-kbp windows across Non-SD (gray), SD (blue), and SD98 (orange), discovered with short-read sequencing (SRS, left) and long-read sequencing (LRS, right) technologies. Number at the bottom represents the total number of 1-kbp windows defined for each region. **(F)** Assessment of precision and recall across eight individuals sequenced with Illumina short-read sequencing and PacBio long-read sequencing reads, for all regions (left) and only accessible sites (right). (**G)** Percentage of short-read accessible bases versus percentage of bases within SD98 regions for 25-kbp windows genome-wide used in Tajima’s D calculations. **(H)** Distribution of Tajima’s D values calculated using 1KGP SNPs from individuals of African ancestry across 25-kbp overlapping protein-coding genes (green), unprocessed pseudogenes (purple), other genes (blue) and no genes (red), in non-duplicated (nonSD) and SD98 regions. *p*-values were calculated using a Mann-Whitney U test. ns: non-significant; * ≤ 0.05, **** ≤ 0.0001. **(I)** Tajima’s D values from individuals of the 1000 Genomes Project were calculated across 25-kbp windows genome-wide (gray) and in SD98 region (orange) divided per superpopulation. Only outlier values in the upper 95th percentile or bottom 5th percentile are shown, plotted across human autosomal chromosomes on the x-axis. Human duplicated genes within windows with outlier D values are highlighted. Ancestries depicted include African (AFR), East Asian (EAS), South Asian (SAS), and American (AMR). **(J)** Assessment of variant association depletion in SD and SD98 regions in short-read-based databases. Included databases: GWAS catalog, ClinVar, and GTEx eQTL. Observed variation is represented in vertical lines for SD (blue) and SD98 (orange) regions, and density plots represent empirical distribution of randomly sampled sites of the same size as SD or SD98 regions. **(K)** Human duplicated genes with significant copy-number differences between autistic probands and unaffected siblings from the Simons Simplex Collection. Significant differences were obtained using a Wilcoxon signed-rank test FDR-adjusted *q*-value < 0.05.

**Figure S2. Human brain expression of duplicated genes related to Figure 2. (A)** Intersection between human duplicated genes expressed (TPM≥1) across prenatal datasets. **(B)** Gene expression across human-duplicated gene subsets in log_2_(TPM) in the CORTECON dataset, spanning pluripotency to upper layer formation, and lymphoblastoid cell line data (n=69) ^66^, stratified by copy number (CN) category. **(C, D)** Module eigengenes from weighted gene co-expression network analysis (WGCNA) of prenatal BrainSpan samples from the prefrontal cortex **(C)** and CORTECON **(D)**. Each module is represented by a randomly assigned color stated above each plot. Numbers in parentheses represent the total number of genes assigned to the module. Stars represent modules enriched on different gene categories, including gene ontology (GO) terms (red), SD98 genes (light blue), human-duplicated genes (dark blue), autism-associated (ASD) genes (yellow), and genomic hotspots from Sattertrom et al. ^191^ (green). Colored bars at the bottom indicate different ages in post-conception weeks (PCW) for BrainSpan and different developmental stages of *ex vivo* neurogenesis for CORTECON. **(E)** Network diagram of the C-yellow module. Only genes within human-duplicated gene families (red), SD98 (pink) and autism-associated (yellow) categories with high module membership are depicted. Genes with asterisks are non-syntenic with the chimpanzee reference (PanTro6) and bold borders are within ±500-kbp of a genomic disorder hotspot.

**Figure S3. Matched neurodevelopment staging of human, mouse, and zebrafish related to Figure 3 and Discussion.** Depicted are principal component analyses of brain single-cell RNA-sequencing samples from **(A)** mouse ^75^ and **(B)** zebrafish ^76^, and (**C**) matched mouse and zebrafish samples to human developmental stages from the BrainSpan dataset.

**Figure S4. Detailed genetic analysis of priority human-duplicated genes related to Figure 4 and STAR Methods. (A)** Sequenced samples used for pHSD variant analysis from **(A)** draft human diploid assemblies included in the Human Pangenome Reference Consortium (HPRC, n=47) and Human Genome Structural Variation Consortium (HGSVC, n=9) and **(B)** capture strategy followed by PacBio HiFi long-read sequencing (cHiFi) from the 1000 Genomes Project (1KGP) and Human Pangenome Reference Consortium (n=200; n=144 unrelated). World maps represent sample sites for each ancestry with counts depicted. **(C)** Benchmarking cHiFi sequencing variants by comparing sequencing coverage between derived and ancestral paralogs (left), unique exons and duplicated exons (middle), and tiled versus untiled regions (right). **(C, D)** Impact of **(C)** mapping quality (MAPQ) and genotyping confidence thresholds, and **(D)** per-sample genotype quality and minimum read depth thresholds on the total number of variant sites (left), variant sites with excess heterozygosity (middle), and median Mendelian concordance across 18 trios (right) using cHiFi reads. **(F)** Heterozygous-site densities across duplicated portions of pHSD captured loci. Variants were identified for HPRC and HGSVC samples (top; n=56) and non-redundant unrelated cHiFi individuals (bottom; n=144). Ancestries depicted include African (AFR), European (EUR), East Asian (EAS), South Asian (SAS), and American (AMR). **(G)** The human genetic variation landscape across *SRGAP2C* locus with 1KGP genome-wide outlier Tajima’s D value (shaded region) as well as and Tajima’s D plots derived from HPRC/HGSVC assembly-derived SNVs using 6-kbp windows and 500-bp steps. **(H, I, J)** Assembly-derived SNVs were also used to characterize nucleotide diversity π (top) and Tajima’s D (bottom) across corresponding duplicated exons, calculated in 15-kbp sliding windows with 1-kbp steps, for human duplicated gene paralogs **(H)** *GPR89*, **(I)** *ROCK1*, and **(J)** *FAM72*. **(K)** Human genetic variation landscape of the *CD8B* locus in T2T-CHM13v1.0 reference genome in the UCSC browser, with HPRC/HGSVC intermediate allele frequency variants, and derived Tajima’s D values calculated in 6-kbp windows with 500-bp steps. Haplotype networks for all HPRC/HGSCV continuous haplotypes in addition to chimpanzee (panTro6) are plotted for each highlighted region, encompassing 6-kbp of sequence. **(L)** Folded Site frequency spectrum with minor allele frequency (MAF) calculated across 35-kbp regions overlapping *CD8B* (light gray) and *CD8B2* (dark gray) from variants detected in the combined dataset including long-read assemblies and capture PacBio HiFi sequencing from individuals of AFR ancestry (n=88 individuals), EUR ancestry (n=29 individuals), and AMR ancestry (n=18 individuals). Three individuals of EUR ancestry (NA20582, NA20525, NA20542) were excluded from this analysis. *p*-values were obtained comparing *CD8B* MAF distribution between populations using Kolmogorov-Smirnov test.

**Figure S5. Analysis of human duplicated priority genes using zebrafish related to Figure 5 and STAR Methods. (A–C)** Endogenous gene expression of pHSD zebrafish orthologs during development. **(A)** Temporal expression between 0 and 120 hours post-fertilization using published data ^106^ of the zebrafish orthologs of the selected pHSDs. Shaded area corresponds to the brain development period in zebrafish embryos ^111^ of the zebrafish orthologs of the selected pHSDs. **(B)** Expression of the selected genes in embryonic or adult tissues (data from ^97^). **(C)** Available expression patterns via *in situ* hybridization in the Zebrafish Information Network (ZFIN) ^98^. **(D)** Detection of human mRNA post-injection in ‘humanized’ zebrafish models using RT-PCR of RNA extracted from 3 dpf injected “humanized” larvae (denoted by white stars) and controls (denoted by black stars) with primers targeting eight human-specific mRNAs (*SRGAP2C*, *ARHGAP11B*, *GPR89B*, *PDZK1P1*, *PTPN20CP*, *NPY4R*, *FAM72B*, *FRMPD2B*). **(E)** Description of the number of cells (n) per zebrafish mutant model used for single-cell transcriptomic analysis.

**Table S1. Genetic variants analysis of SD98 genes related to Figure 1, Discussion, and STAR Methods. (A)** SD-98 genes (>1 exon overlapping segmental duplications with over 98% identity) in T2T-CHM13v1.0, database intersections, and brain RNA-seq expression (TPM: transcripts per million). **(B)** SD-98 genes in chromosomes T2T-CHM13 (v1.0) X and T2T-HG002Y (hs1). **(C)** SD98 gene clustering into gene families based on shared exons and similar famCN (MAD<1) between paralogs. Copy number was calculated only for protein coding and unprocessed pseudogenes, but other overlapping gene features (i.e. lncRNA) were reported. **(D)** Predicted evolutionary status of SD98 gene families. **(E)** Copy-number variation analysis in SD98 regions. parCN: paralog-specific copy-number. **(F)** Outlier Tajima’s D values across SD98 windows (>10% SD98) at least 50% accessible and carrying 5 or more SNPs. **(G)** *De novo* copy-number events of SD98 genes identified in the SSC.

**Table S2. Human brain expression of SD98 genes related to Figure 2. (A)** BrainSpan WGCNA module assignment. **(B)** Gene ontology overrepresentation of BrainSpan WGCNA modules. **(C)** Cortecon WGCNA gene-module assignment. **(D)** Gene family co-expression concordance using CORTECON WGCNA. **(E)** Gene ontology overrepresentation of CORTECON WGCNA modules.

**Table S3. Modeling SD98 genes in mouse and zebrafish related to Figure 3 and Discussion. (A)** Mouse and zebrafish orthologs of SD98 gene families. **(B)** SD98 genes fetal brain expression in human, mouse and zebrafish orthologs.

**Table S4. Sequence and variant analysis of priority human-specific duplicated (pHSD) genes related to Figure 4 and STAR Methods. (A)** Summary of selected pHSD genes, canonical transcripts, variant calling, and variant effect prediction. **(B)** Individuals sequenced with capture PacBio HiFi long read sequencing. **(C)** Coordinates of captured pHSD regions and summary of long-read sequencing variant discovery. **(D)** Oligonucleotide baits design for cHiFi sequencing. **(E)** Summary statistics of cHiFi sequencing. **(F)** Variant effect prediction across pHSD paralogs. **(G)** pHSD coding variants and allele frequencies. **(H)** Ka/Ks, biallelic SNPs, pN/pS, and Direction of Selection across pHSD paralogs. **(I)** Comparison of dN/dS under different models using codeml.

**Table S5. Details of zebrafish models of priority human-duplicated (pHSD) genes related to Figure 5, Discussion, and STAR Methods. (A)** Oligonucleotide sequences used in this study. **(B)** Survival of mutant zebrafish larvae. **(C)** Distribution of the 3,146 images of larvae for morphological assessments. **(D)** Raw morphometric data for all zebrafish models of the selected pHSD genes for functional characterizations. **(E)** Statistical results from the morphological comparisons across zebrafish models of the selected pHSD genes.. **(F)** Description of the sci-RNA-seq identified clusters from heads of 3 dpf zebrafish larvae. **(G)** Marker genes for all identified clusters in the sci-RNA-seq data from heads of 3 dpf zebrafish larvae. **(H)** Gene ontology (GO) terms enriched in DEGs across zebrafish models of the selected pHSD genes for forebrain and midbrain. **(I)** GO terms enriched in DEGs for *SRGAP2* mutant zebrafish models. **(J)** GO terms enriched in DEGs for *ARHGAP11B* humanized zebrafish model.

**Table S6. Zebrafish mutant models *gpr89* and *frmpd2* related to Figure 6 and STAR Methods. (A)** Differentially expressed genes (DEGs) between *GPR89B* and *gpr89* knockout (KO) models and their respective controls. **(B)** Gene ontology (GO) terms enriched in DEGs between *GPR89B* and *gpr89* KO models and their respective controls. **(C)** DEGs between *FRMPD2*B and *frmpd2* KO models and their respective controls. **(D)** GO terms enriched in DEGs between *FRMPD2B* and *frmpd2* KO models and their respective controls. **(E)** Stable mutant zebrafish alleles.

**Data S1. Weighted gene co-expression analysis of human brain samples related to Figure 2 and STAR Methods.**

