## Supplemental Figures & Data for "Human-specific gene expansions contribute to brain evolution": DataS1.pdf

### Data S1: Weighted gene co-expression analysis of human brain samples related to Figure 2 and STAR Methods.

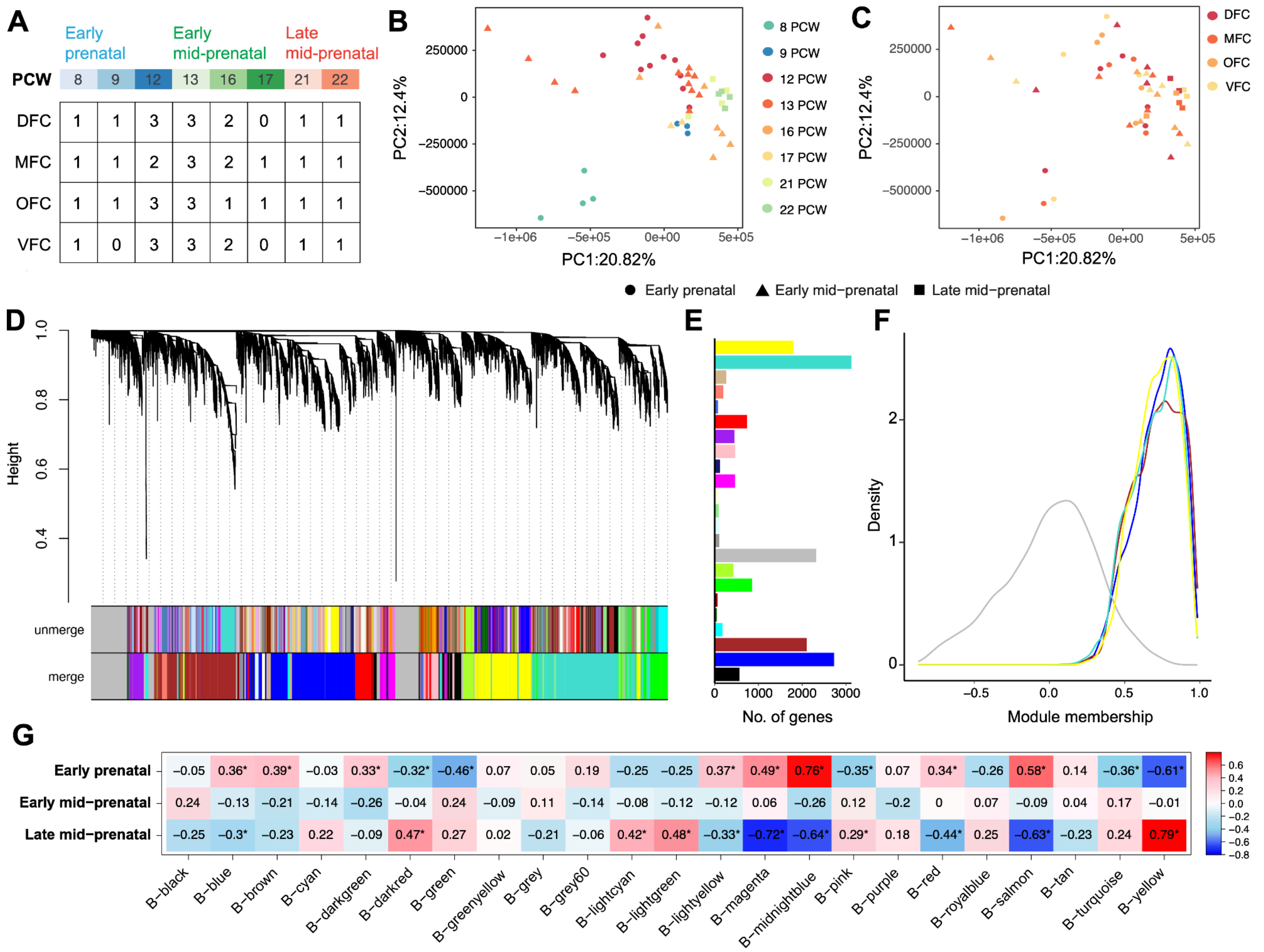


**DataS1-Figure 1. BrainSpan data set pre-processing and clustering using gene co-expression network analysis.** (**A**) BrainSpan dataset sample numbers classified by weeks post-conception (PCW), developmental epoch and brain region (DFC: dorsolateral prefrontal cortex, VFC: ventrolateral prefrontal cortex, MFC: medial prefrontal cortex, OFC: orbital prefrontal cortex). (**B**) PCA of BrainSpan samples by developmental epoch and age in weeks post-conception. (**C**) PCA of BrainSpan samples by developmental epoch and brain region. (**D**) Hierarchical clustering of genes for WGCNA module assignment. Modules are shown before and after merging highly correlated modules. (**E**) Number of genes assigned to co-expression modules annotated with random colors. (**F**) Distribution of module membership across genes within B-turquoise, B-blue, B-brown, B-yellow, and B-grey modules. (**G**) Modules eigengenes association with developmental epochs. Numbers inside boxes and their colors indicate Pearson correlation values. Significant correlation (p-value<0.05) are highlighted with an asterisk.

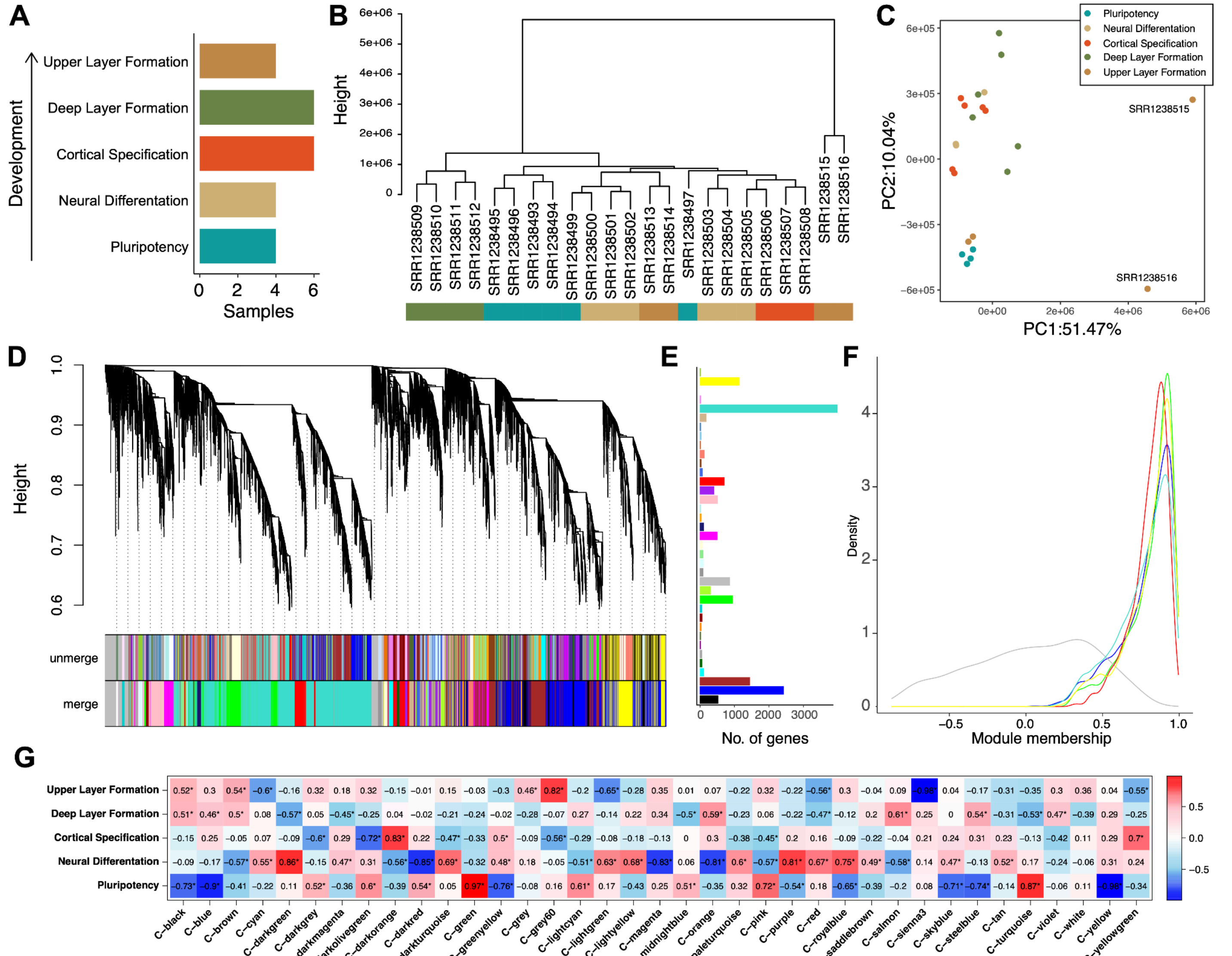


**Data S1-Figure 2. CORTECON data set pre-processing and clustering using gene co-expression network analysis.** (**A**) CORTECON dataset sample numbers classified by developmental stage. (**B**) Hierarchical clustering of CORTECON samples. (**C**) PCA of CORTECON samples. (**D**) Hierarchical clustering of genes for WGCNA module assignment. Modules are shown before and after merging highly correlated modules. (**E**) Number of genes assigned to co-expression modules annotated with random colors. (**F**) Distribution of module membership across genes within C-turquoise, C-blue, C-yellow, C-green, C-red, and C-grey modules. (**G**) Modules eigengenes association with developmental stages. Numbers inside boxes and their colors indicate Pearson correlation values. Significant correlation (p-value<0.05) are highlighted with an asterisk.
