## Supplementary figures and images for "Human-specific gene expansions contribute to brain evolution"

### FigS1.png

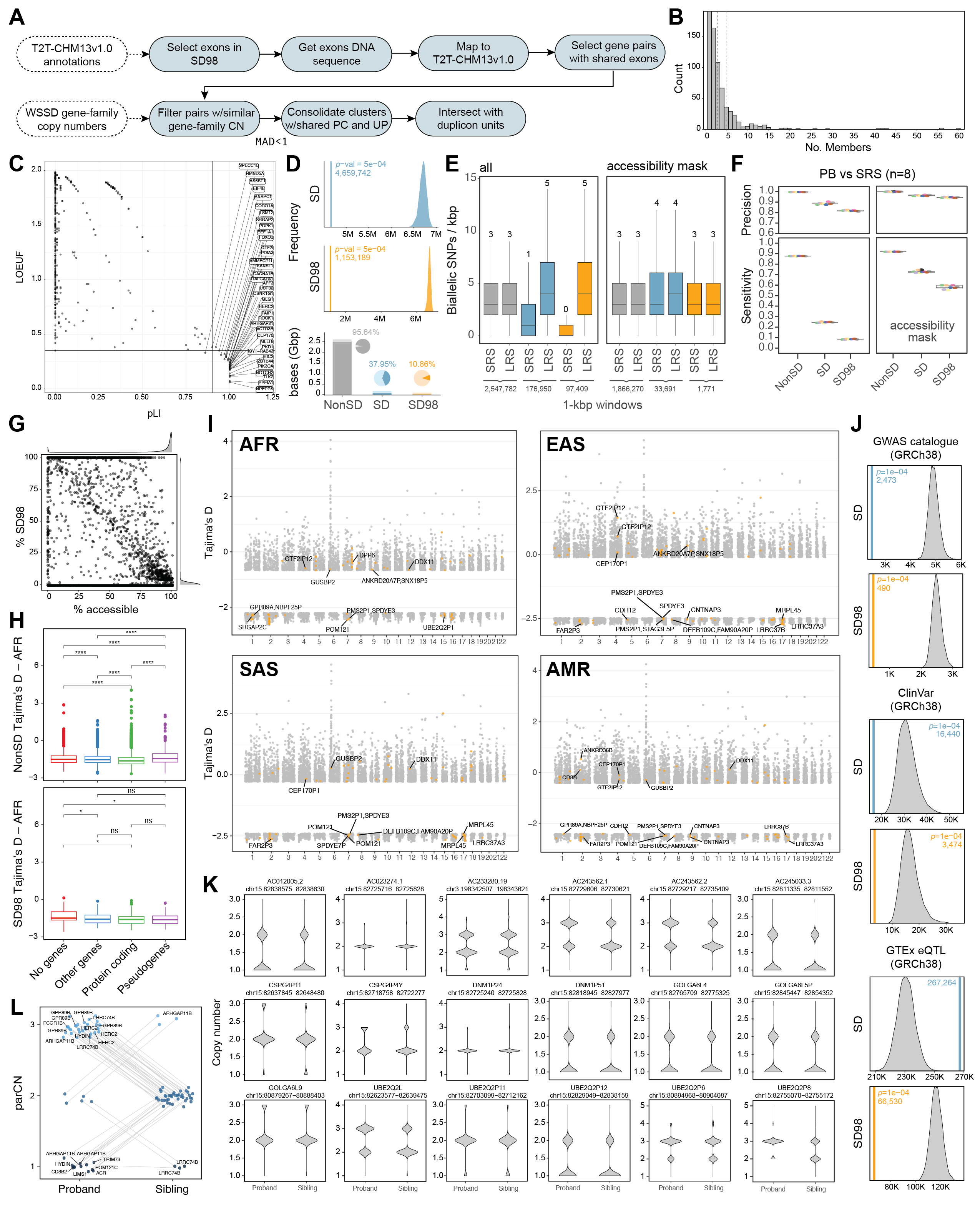

### FigS2.png

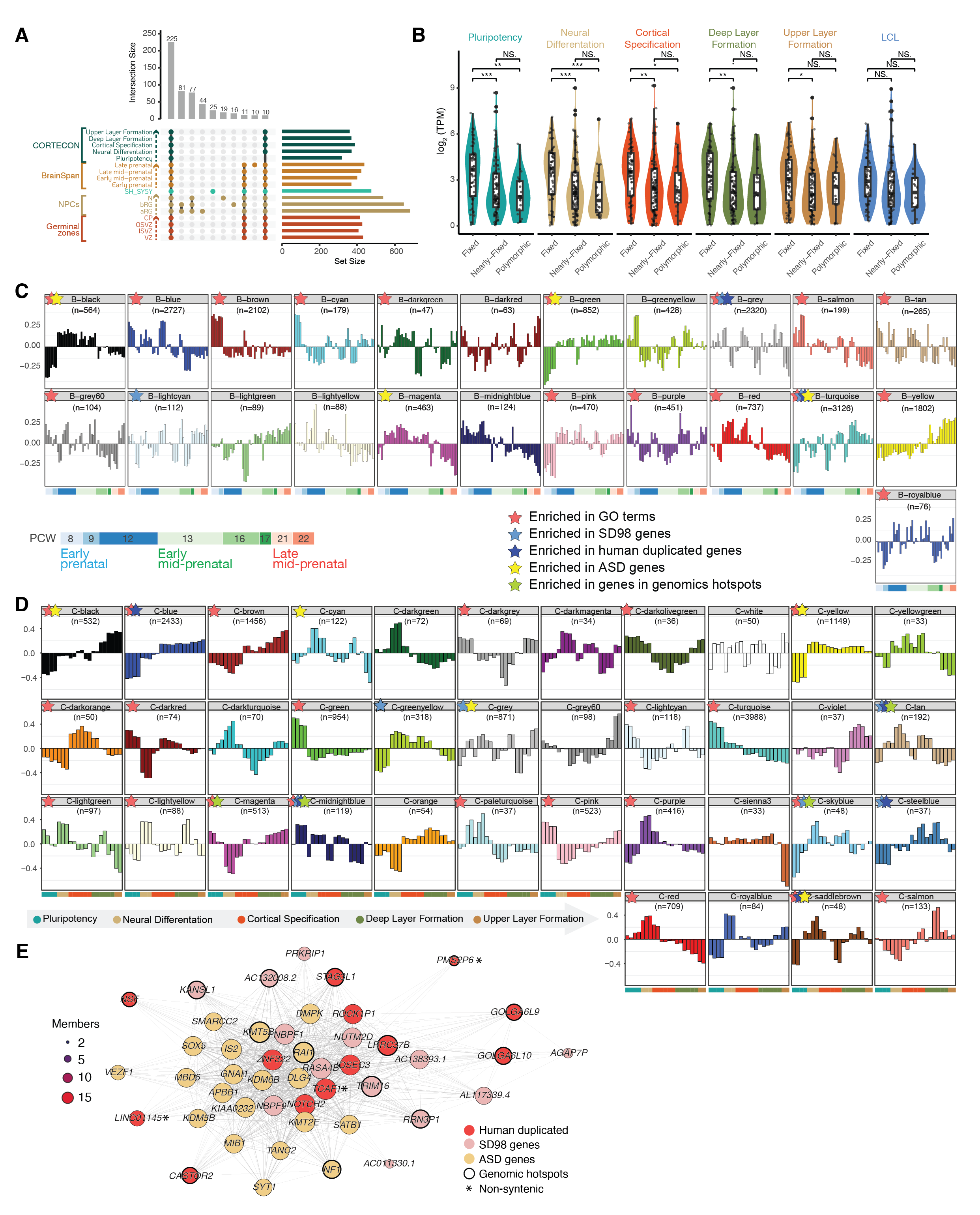

### FigS3.png

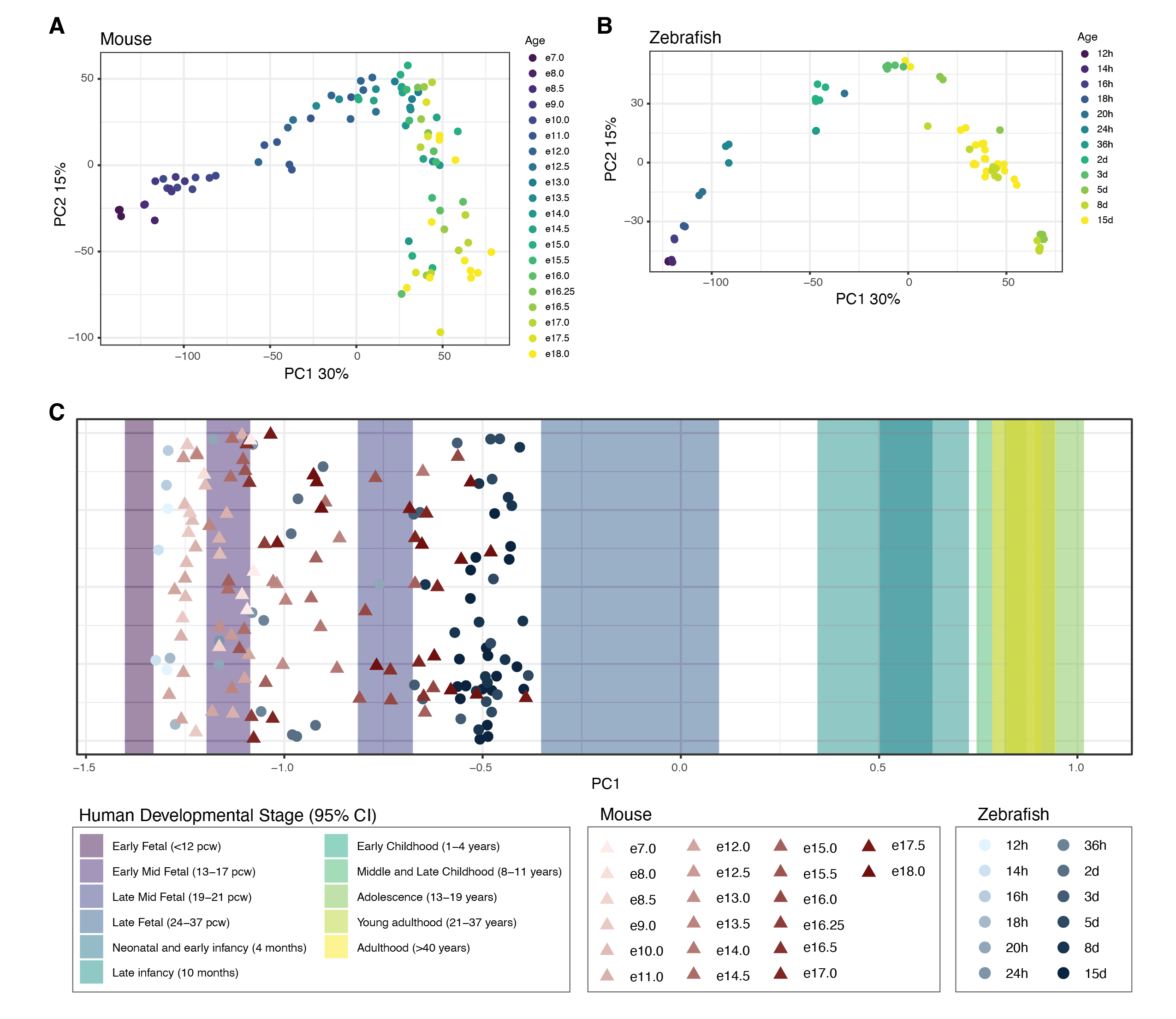

### FigS4.png

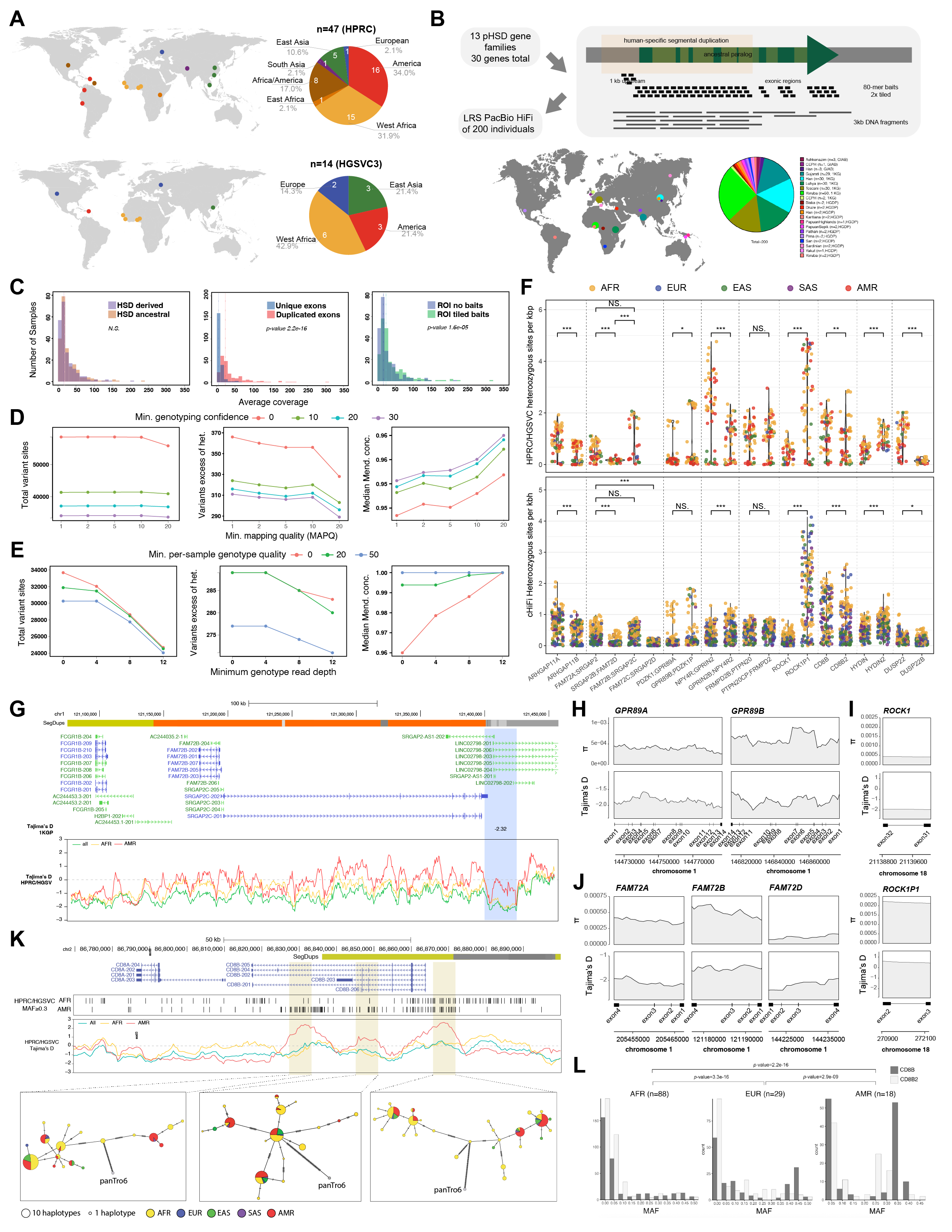

### FigS5.png

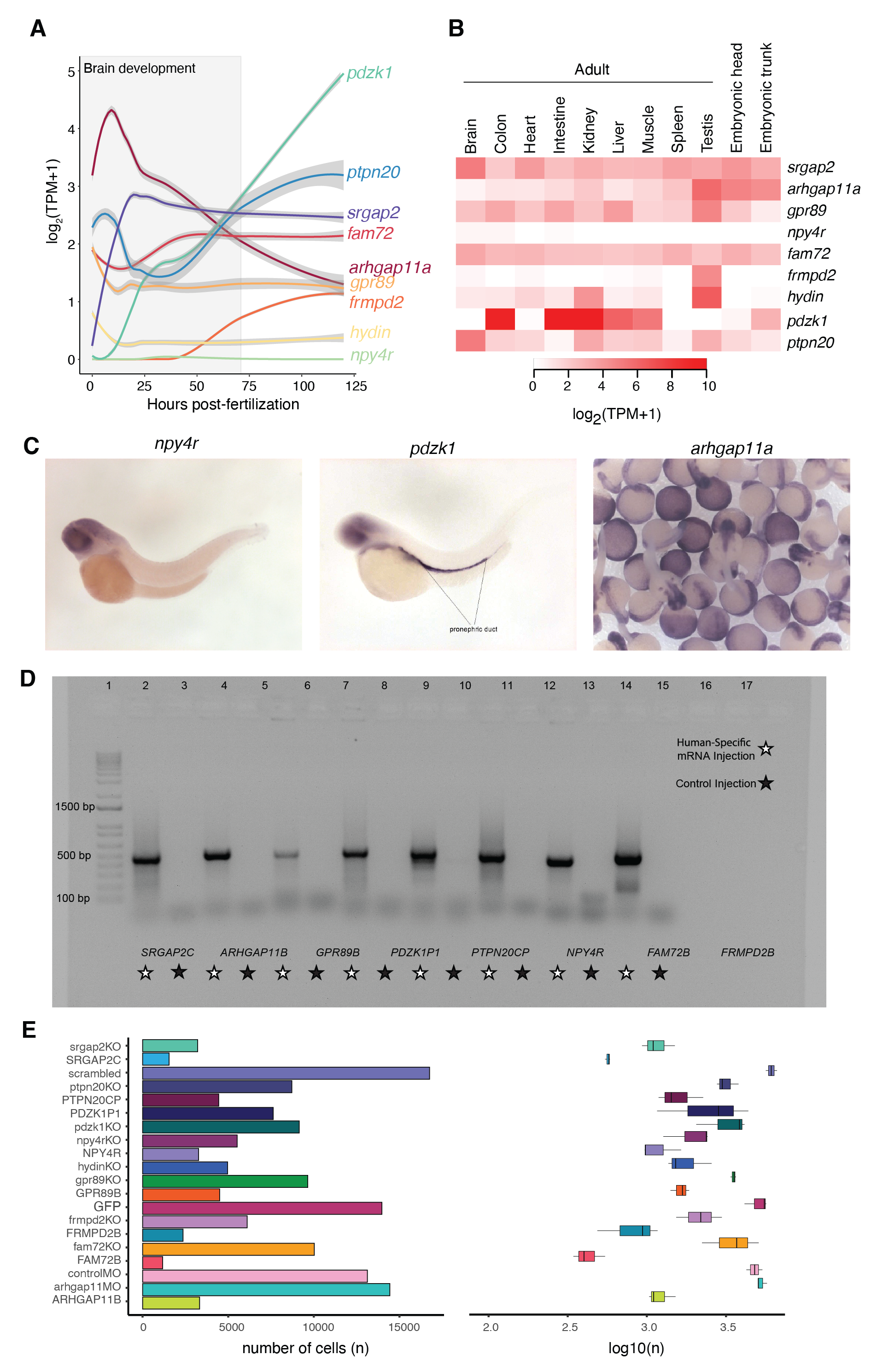
